# Chromatin dynamics during hematopoiesis reveal discrete regulatory modules instructing differentiation

**DOI:** 10.1101/2020.04.02.022566

**Authors:** Grigorios Georgolopoulos, Nikoletta Psatha, Mineo Iwata, Andrew Nishida, Tannishtha Som, Minas Yiangou, John A. Stamatoyannopoulos, Jeff Vierstra

**Author notes:** Correspondences to and/or. These authors contributed equally (Grigorios Georgolopoulos, Nikoletta Psatha). Department of Development and Regeneration, KU Leuven, Leuven, Belgium.

## Abstract

Lineage commitment and differentiation is driven by the concerted action of master transcriptional regulators at their target chromatin sites. Multiple efforts have characterized the key transcription factors (TFs) that determine the various hematopoietic lineages. However, the temporal interactions between individual TFs and their chromatin targets during differentiation and how these interactions dictate lineage commitment remains poorly understood. We performed dense, daily, temporal profiling of chromatin accessibility (DNase I-seq) and gene expression changes (total RNA-seq) along *ex vivo* human erythropoiesis to comprehensively define developmentally regulated DNase I hypersensitive sites (DHSs) and transcripts. We link both distal DHSs to their target gene promoters and individual TFs to their target DHSs, revealing that the regulatory landscape is organized in distinct sequential regulatory modules that regulate lineage restriction and maturation. Finally, direct comparison of transcriptional dynamics (bulk and single-cell) and lineage potential between erythropoiesis and megakaryopoiesis uncovers differential fate commitment dynamics between the two lineages as they exit pluripotency. Collectively, these data provide novel insights into the global regulatory landscape during hematopoiesis.

## Introduction

The temporal activation of stage-specific regulatory DNA instructs lineage specific gene expression programs that underpin cellular fate and potential. The establishment and maintenance of regulatory DNA is mediated by the combinatorial engagement of sequence-specific transcription factors (TFs) that bind in the place of a canonical nucleosome. Over the course of cellular differentiation programmed shifts in the global transcription factor milieu drive extensive re-organization of chromatin^1,2^, where silencing of regulatory DNA associated with alternate lineage and the *de novo* activation of lineage-restricted elements result in the narrowing of the epigenetic and functional landscape^3^. However, it is unclear how and when regulatory DNA is dynamically activated and silenced during cell state transitions to establish lineage restricted gene expression programs and how these epigenetic changes relate to developmental potential.

Hematopoiesis is a prototypical system to study how genetically and epigenetically encoded programs are established during cellular differentiation^4–6^. Conventionally, hematopoiesis is depicted as a discrete hierarchical process where a multipotent hematopoietic stem and progenitor cell (HSPC) traverses a sequence of bifurcating decisions, mediated by the expression of lineage-specific TFs, with each decision resulting in an increasingly restricted fate potential. Historically, the characterization of the gene regulatory programs involved in the transition from HSPCs to terminal fates has relied on the identification of differential transcriptional programs from isolated discrete populations using defined cell surface markers^7–10^. While this approach has led to the identification of master regulatory transcription factors^10,11^ that define many of the major hematopoietic cell lineages and has enabled a systematic mapping of their steady-state regulatory landscapes^9,12^, interrogation of discretely defined populations cannot elucidate the dynamic regulatory events that mark cell-state transitions.

Recently, single-cell chromatin and transcriptional profiling assays have attempted to resolve the spatio-temporal *cis*- and *trans*-dynamics in different stages of hematopoiesis^13–16^. These studies have largely relied on the analysis of either bulk or immunophenotypically isolated populations of steady-state peripheral blood or bone marrow derived cells, whereby hierarchical relationships and developmental trajectories between cell states are predicted computationally. While such experimental approaches have aided in defining major subpopulations of hematopoietic cells and their respective epigenetic and transcriptional landscapes, definition of developmental trajectories within individual lineages from population snapshots is challenging due to the limited sensitivity and the resulting technical and analytical artifacts associated with single-cell genomic assays^17,18^. Additionally, because developmental trajectories are predicted *in silico*, direct association of functional changes (i.e., lineage potential) to intermediate cellular states is not possible^19^.

In order to investigate the dynamics of regulatory and functional events during differentiation, we use human erythropoiesis as a proxy for hematopoietic development. The transition from HSPCs to terminally differentiated enucleated red blood cells involves a series of morphologically, functionally, and phenotypically distinguishable states. Multiple efforts relying on the isolation of these states have exhaustively characterized key transcriptional regulators^20,21^ and chromatin elements implicated in erythropoiesis^9,22^. However, a general understanding of the temporal interplay between individual *cis*- and *trans*-elements and how these establish stage-specific transcriptional programs and lineage commitment during hematopoiesis remains rudimentary. Furthermore, because erythrocytes share their developmental origins of with other myeloid lineages (granulocytic/monocytic and megakaryocytic), erythropoiesis represents an ideal system to study how lineage choice is genetically and epigenetically encoded.

Here, we capitalize on the *ex vivo* human differentiation scheme where dense unbiased sampling of the populations captures the dynamics of chromatin accessibility and gene expression during differentiation with a completely defined developmental trajectory. DNase I-seq and gene expression profiling (bulk and single cell) time-course during erythropoiesis coupled with lineage potential assays and morphological characterization, enabled the assignment of distal elements (alone or in combination) to target genes and individual TFs to their target DHSs which collectively comprise discrete regulatory modules associated with lineage potential. Comparing the activity patterns of the TF regulatory modules in the erythroid lineage to the closely related megakaryocytic lineage, provides insights into how these modules instruct lineage commitment. Collectively, our findings provide key insights into the organization of the functional epigenetic landscape during hematopoietic differentiation and its relation to lineage-potential.

### Dense mapping of the temporal dynamics of *cis*- and *trans*-elements during erythropoiesis

Human erythropoiesis was induced *ex vivo* for 12 days using an established differentiation protocol^23^ that faithfully recapitulates the major features of *in vivo* erythropoiesis. Starting from human adult-derived mobilized peripheral blood CD34^+^-enriched HSPCs from 3 healthy donors we cultured the cells in defined media for 12 days (**Figure 1a** and **Methods**). Characteristic features of developing erythroblast cells were confirmed by immunophenotyping using canonical cell-surface markers of early (CD117, C-Kit) and late (CD235a, Glycophorin A) erythropoiesis as well as morphologically by hematoxylin-eosin staining of cell smears (**Supplementary Figure 1**).

**Figure 1.**
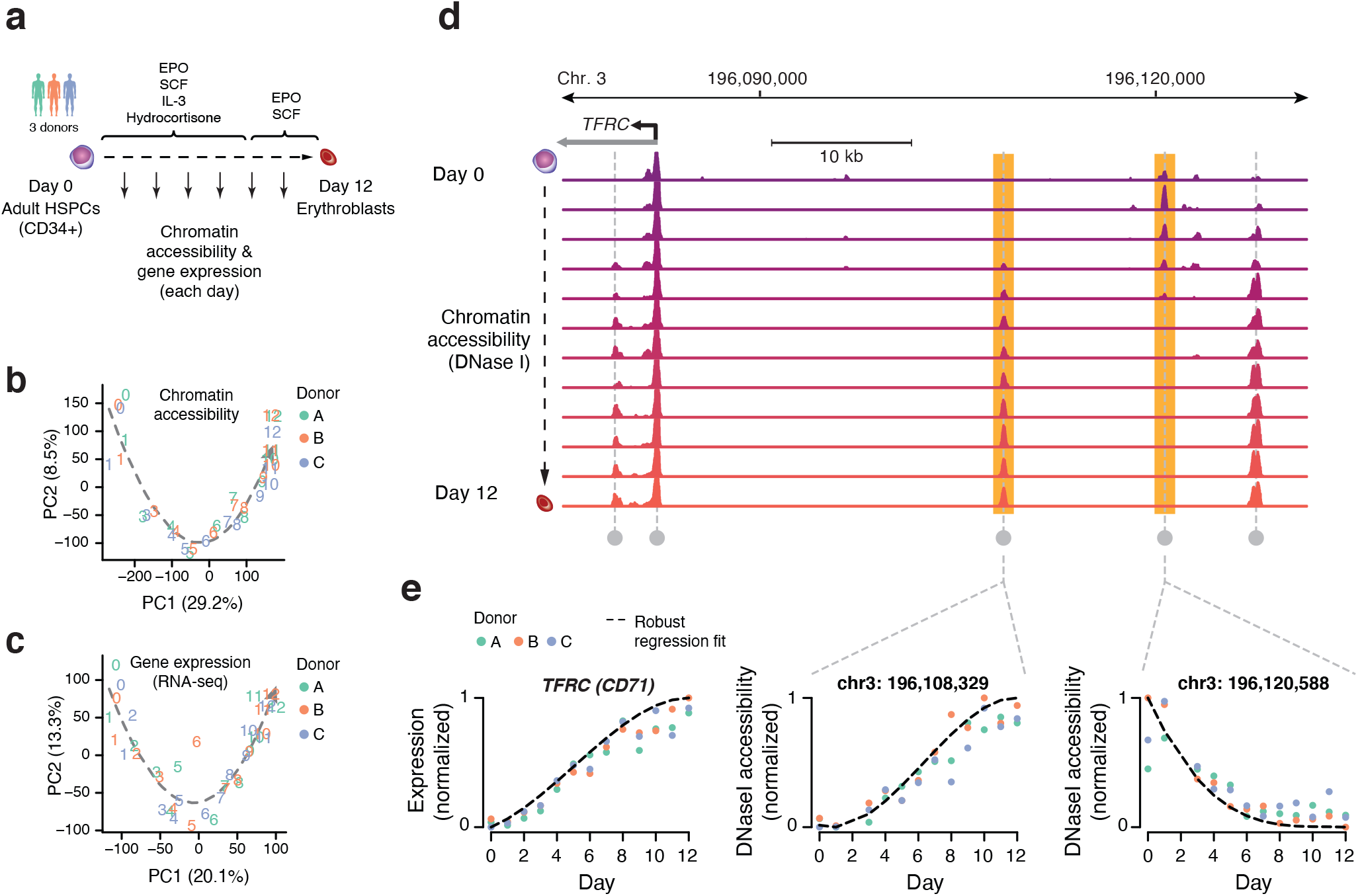
Comprehensive identification of regulatory landscape developmental dynamics. (a) Dense DNase I-seq and RNA-seq time course with daily sampling during the 12-day *ex vivo* erythroid differentiation induced from CD34^+^ HSPCs. (b) PCA analysis using all detected DHSs (79,085 Hotspots 5% FDR) across all samples (12 time points, 3 donors). The arrow denotes the differentiation trajectory from day 0 to day 12. (c) PCA analysis using all 24,849 detected genes across all samples (13 time points, 3 donors). The arrow denotes the differentiation trajectory from day 0 to day 12. (d) Chromatin accessibility tracks for each day of differentiation with DNase Hypersensitive Sites (DHSs) harbored around the *TFRC* locus. (e) Identification of significantly changing DHS and genes with robust linear regression analysis. Scatterplots show *TFRC* expression and DNase I density for two upstream DHS. Dots represent normalized values for each of the 3 donors. Dashed line represents the fitted regression spline.

To densely map both chromatin accessibility and transcriptional dynamics during the transition from HSPCs to committed erythroblasts, we subsampled a single continuous culture each day (12 days) and performed DNase I-seq analysis and total RNA-seq (**Figure 1a,b**). Biological replicates from CD34^+^ HSPCs from 3 donors were highly reproducible for both chromatin accessibility and gene expression profiles where the majority of the observed variability was accounted for by developmental trajectory (i.e., sampling days) (**Figure 1b,c** and **Supplementary Figure 2a**), as biological replicates were highly correlated (**Supplementary Figure 2b,c**). For many individual DHSs and genes, we observed quantitative changes in chromatin accessibility and expression over the course of differentiation highlighted by quantitative trajectories of opening or closing (**Figure 1d,e**). Notably, accessibility changes were mostly confined to compact regions of the genome (∼200bp average DHS width). In many cases, we observed both opening and closing events within close proximity (**Figure 1d**), indicating focal regulation^24^ of chromatin structure in contrast to previous reports that chromatin changes during differentiation occur over large domains^1,25^.

**Figure 2.**
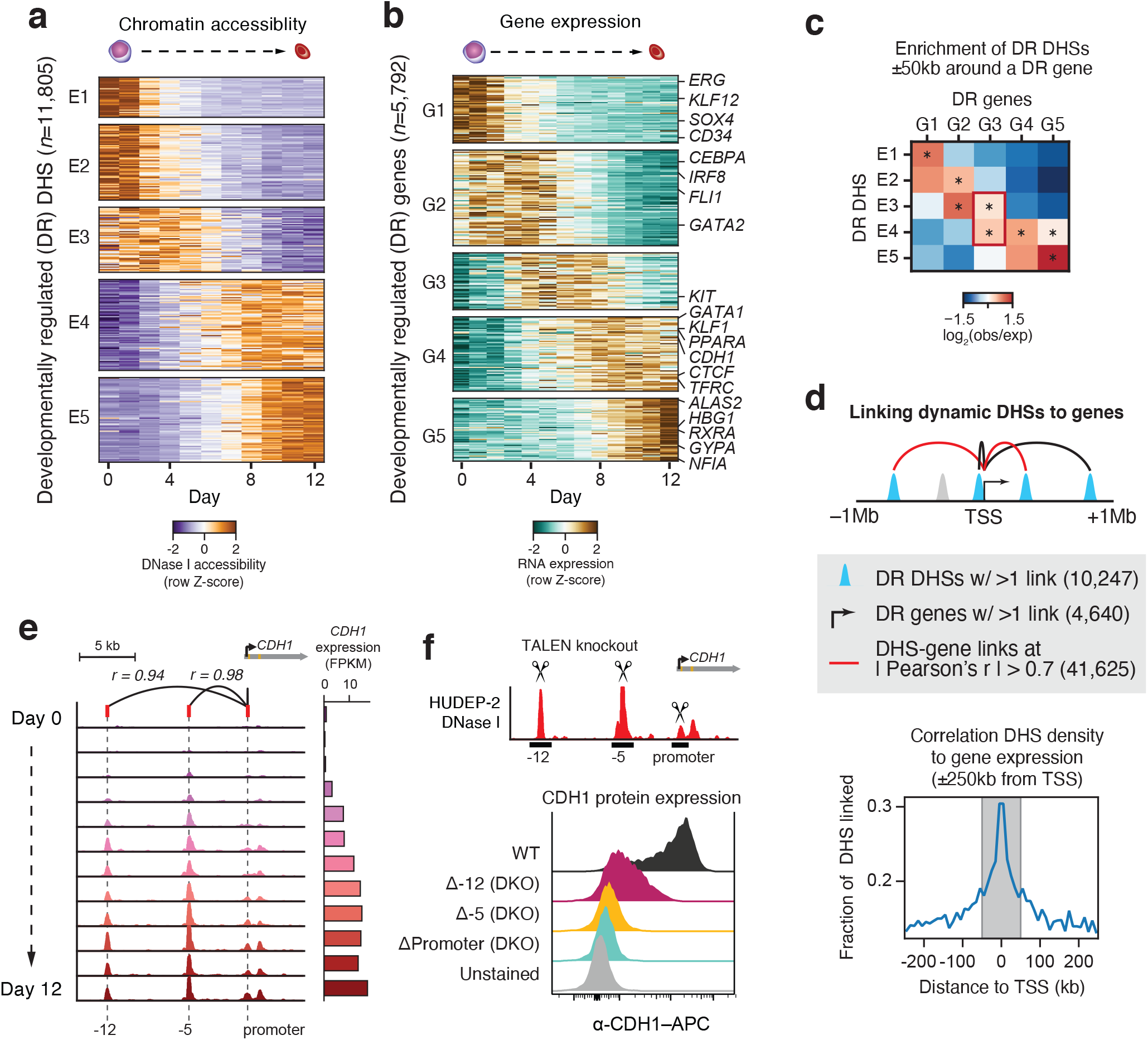
Temporal compartmentalization of the e *cis-* and *trans*- epigenetic landscape during erythropoiesis exhibits. (a) *K*-means clustering of 11,805 changing DHS resulted in 5 clusters (E1-E5) with sequential activity profile for each cluster. Values are z-score of per day average normalized DHS counts from 3 donors. (b) *K*-means clustering of 5,792 developmentally regulated genes resulted in 5 clusters (G1-G5). Values are z-score of per day average normalized FPKM from 3 donors. (c) A matrix showing the enrichment score (log2-ratio observed over expected) for any given DHS cluster, around each developmentally regulated DHS (±50kb from TSS). Highlighted in red is cluster G3 which is enriched for both late downregulated DHS of cluster E3 and early upregulated from cluster E4. ^*^ *X*^2^ test *p*-value < 0.05 (d) Correlation density plot between developmental genes and developmental DHS ±250kb around the gene promoter. Grey shaded area highlights enrichment of correlations within ±50kb around the gene promoter. (e) DNase I accessibility track of the *CDH1* locus during erythroid differentiation, highlighting the accessibility of 3 nearby DHS correlated to *CDH1* expression. (f) DNase I accessibility track of the *CDH1* locus in HUDEP-2 cells depicting the genetic knockout of the *CDH1* promoter and two upstream DHS (−12, and - 5) (above) along with the resulted ablation in CDH1 protein expression as assessed by flow cytometry (below).

To systematically identify developmentally responsive *cis*-elements, we leveraged the observed continuity of DHS signal over adjacent days (**Figure 1d**) and modelled DNase I cleavage density against differentiation time-points (**Methods**). We determined significance by comparing our full model to a reduced model (intercept-only; not accounting for developmental time) and performing a likelihood ratio test (**Methods**). Of the total 79,085 DHSs accessible in 2 or more samples/replicates, we conservatively identified 11,805 (14.9%) significantly changing DHSs (adjusted *p* < 10^−5^ and fold-change >2), nearly evenly grouped between activated and silenced (45% and 55%, respectively) (Supplementary Table 1). A similar analytical approach applied to the RNA expression data identified 5,769 developmentally regulated genes (adjusted *p* < 10^−5^ and fold-change > 2), of which 62% up-regulated and 38% down-regulated over the course of differentiation (**Supplementary Table 2**). Collectively, these data define a high-resolution and quantitative map of chromatin and gene expression dynamics during erythroid differentiation.

### Stage-specific compartmentalization of the *cis*- and *trans*-landscape

PCA indicated that days 5-6 were associated with a critical developmental inflection point during *ex vivo* differentiation (**Figure 1b,c**). We therefore sought to characterize the relationship between temporal chromatin and gene expression dynamics with regards to the observed immunophenotypic and morphological changes present in the population of differentiating cells. We performed unsupervised clustering (*K*-means; *k*=5) on dynamically changing DHSs and developmentally responsive transcripts (**Figure 2a,b**). This analysis revealed a stark partitioning of activated and silenced genes and DHSs into non-overlapping sets that closely paralleled canonical developmental features of erythropoiesis. Particularly, DHSs rapidly silenced within the first days of differentiation (clusters E1 and E2) were found to preferentially harbor binding sequences utilized by the known HSPC regulators such as (HOXA9^26^, RUNX^27^ and ERG^28^) (**Supplementary Figure 3**). Similarly, immediately downregulated transcripts upon induction of differentiation (cluster G1) include these transcription factors as well as structural genes characteristic of CD34^+^ HSPCs (**Figure 2b**). Consistent with PCA (**Figure 1b,c**), a rapid and marked turnover of chromatin and gene expression landscape is observed between days 5-7 where an early erythroid signature appears in both activated DHSs and gene expression, marked by the upregulation of *GATA1, KLF1, PPARA* and *TFRC* (cluster G4). Markers of mature erythropoiesis emerge later in the differentiation (after day 8; cluster G5) with the upregulation of hemoglobins, glycophorin A (*GYPA*) and *ALAS2* (**Figure 2b**). Beyond the temporal partitioning of developmentally regulated DHS and transcripts we observed topological segregation of co-regulated elements. Mapping changing DHS and genes to TADs called from CD34+ HSPCs^29^ and day 11 *ex vivo* differentiated erythroid progenitors^30^ Hi-C data revealed enrichment of individual TADs for stage-specific elements (**Supplementary Figure 3**). Additionally, this partitioning appears more contrasted in late erythroid TADs compared to CD34+, suggesting the establishment of a defined erythroid regulatory landscape.

**Figure 3.**
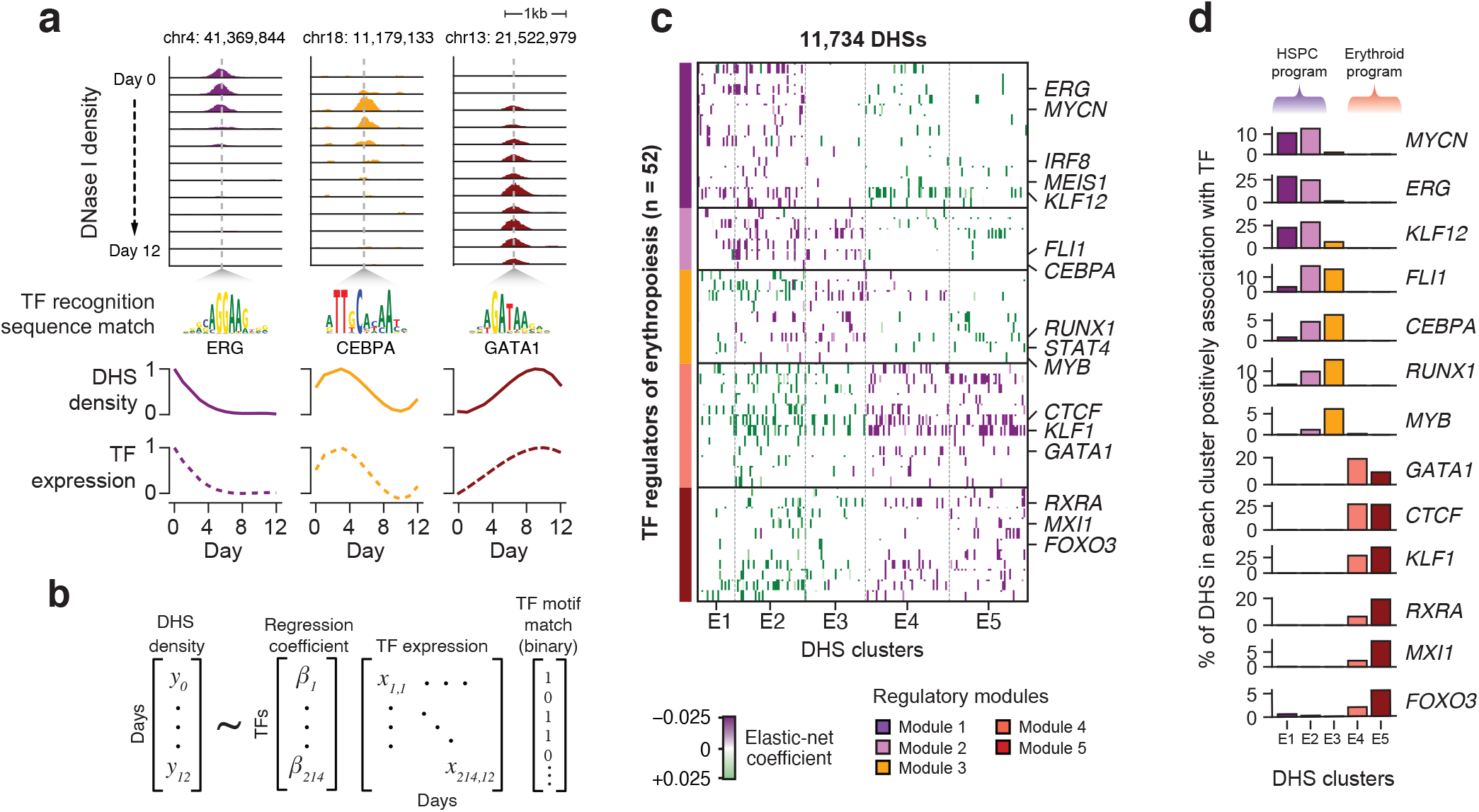
Systematic modelling of *cis*- and *trans*- element temporal interactions reveals discrete regulatory modules during erythropoiesis. (a) Developmental responses of DHS accessibility and transcription factor expression levels were found to be correlated across the genome. (b) The density of a given developmental DHS is modelled after the TF binding motif composition and the expression of the binding TFs using elastic-net regression. The model returns a coefficient for each pair of DHS and binding TF which denotes how strongly (positively or negatively) the TF expression is associated with the accessibility of the particular DHS. (c) Hierarchical clustering of 52 highly connected TFs based on the cosine distances of the regression coefficient from 11,734 DHS reveals 5 clusters of developmentally regulated TFs. Transcription factors along with their positively associated DHS comprise a regulatory module (modules 1-5). (d) The fraction of DHS per cluster positively associated with a TF identifies the major drivers of chromatin accessibility during erythropoiesis.

In addition to canonical activation and downregulation patterns observed, we found a subset of genes exhibiting reproducible transient upregulation (clusters G2 and G3) occurring prior to establishment of the erythroid signature (**Figure 2b**). Transiently upregulated genes are found enriched in transcripts representing myeloid lineages including several myeloid markers (e.g. *MPO, KIT*) as well as the myeloid-specific transcription factor CEBPA. Compatible with gene expression, late closing DHSs in cluster E2 and E3 were enriched in CEBPA recognition sequences (**Supplementary Figure 3**). Moreover, the majority (∼80%) of DHSs in cluster E2 and E3 were found overlapping with DHSs active in other myeloid cell types (macrophages and monocytes) (**Supplementary Figure 5**), denoting a transient emergence of myeloid-related regulatory program prior to erythroid commitment.

**Figure 4.**
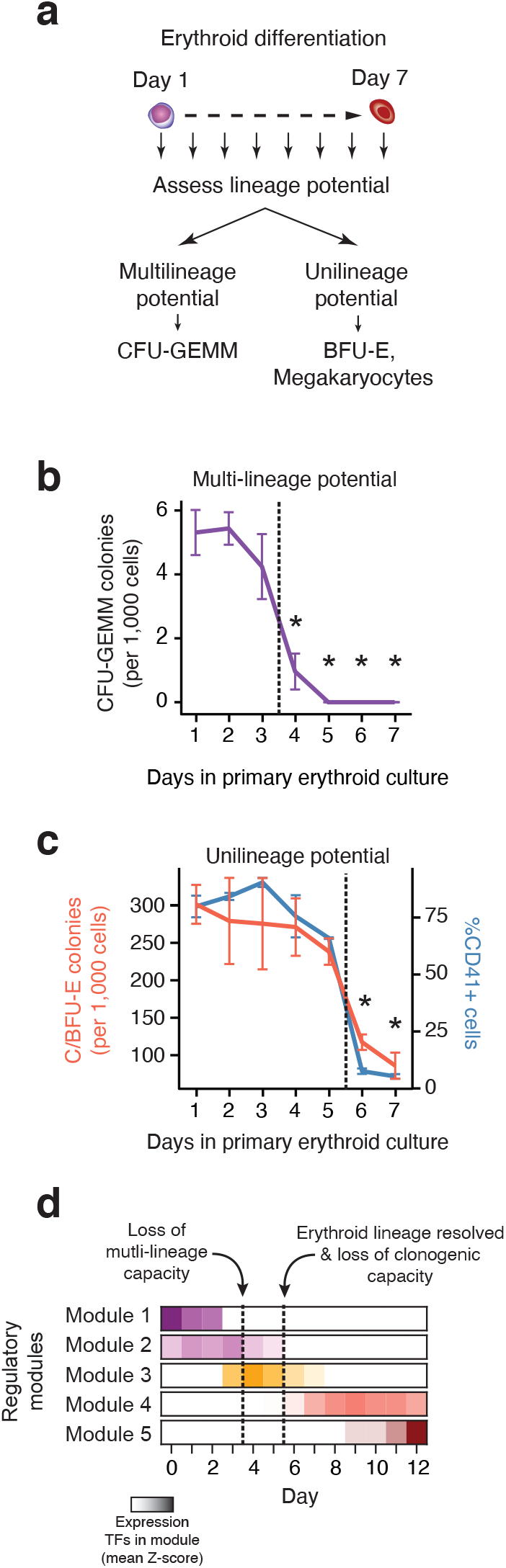
Lineage restriction events during erythropoiesis reflect the sequence of regulatory programs. (a) Schematic diagram of lineage potential assays during the first 7 days of *ex vivo* erythropoiesis. Cells were sampled daily and transferred to lineage-permissive media. Multilineage capacity was determined as frequency of CFU-GEMM progenitors. Erythroid potential as frequency of BFU-Es and megakaryocytic potential as frequency of CD41^+^ cells. (b) Frequency of multipotent CFU-GEMM in methylcellulose assay from cells sampled over the course of erythroid differentiation. (c) Frequency of unipotent erythroid progenitor colonies (BFU-E) in methylcellulose assay (red line) and frequency of CD41^+^ cells after transplantation in secondary megakaryocytic media (blue line) (d) Changes in lineage potential coincide with the transitions between the regulatory modules identified earlier. Transition from modules 1 and 2 to module 3 reflects the loss of multipotency occurring between days 3 and 4, while transition from module 3 to erythroid modules 4 and 5 coincides with the depletion of unipotent progenitors and entry to erythroid maturation (days 5 to 6). Error bars denote ±1 SE of the mean from 4 replicates for colony-forming assays, 2 replicates for CD41^+^ frequency. Asterisk denotes statistically significant difference in CFU-GEMM and BFU-E counts from day 1 (*P*-value < 0.05 Student’s T-test). CFU-GEMM: Colony Forming Unit - Granulocytic, Erythroid, Macrophage, Megakaryocytic. BFU-E: Burst Forming Unit-Erythroid.

**Figure 5.**
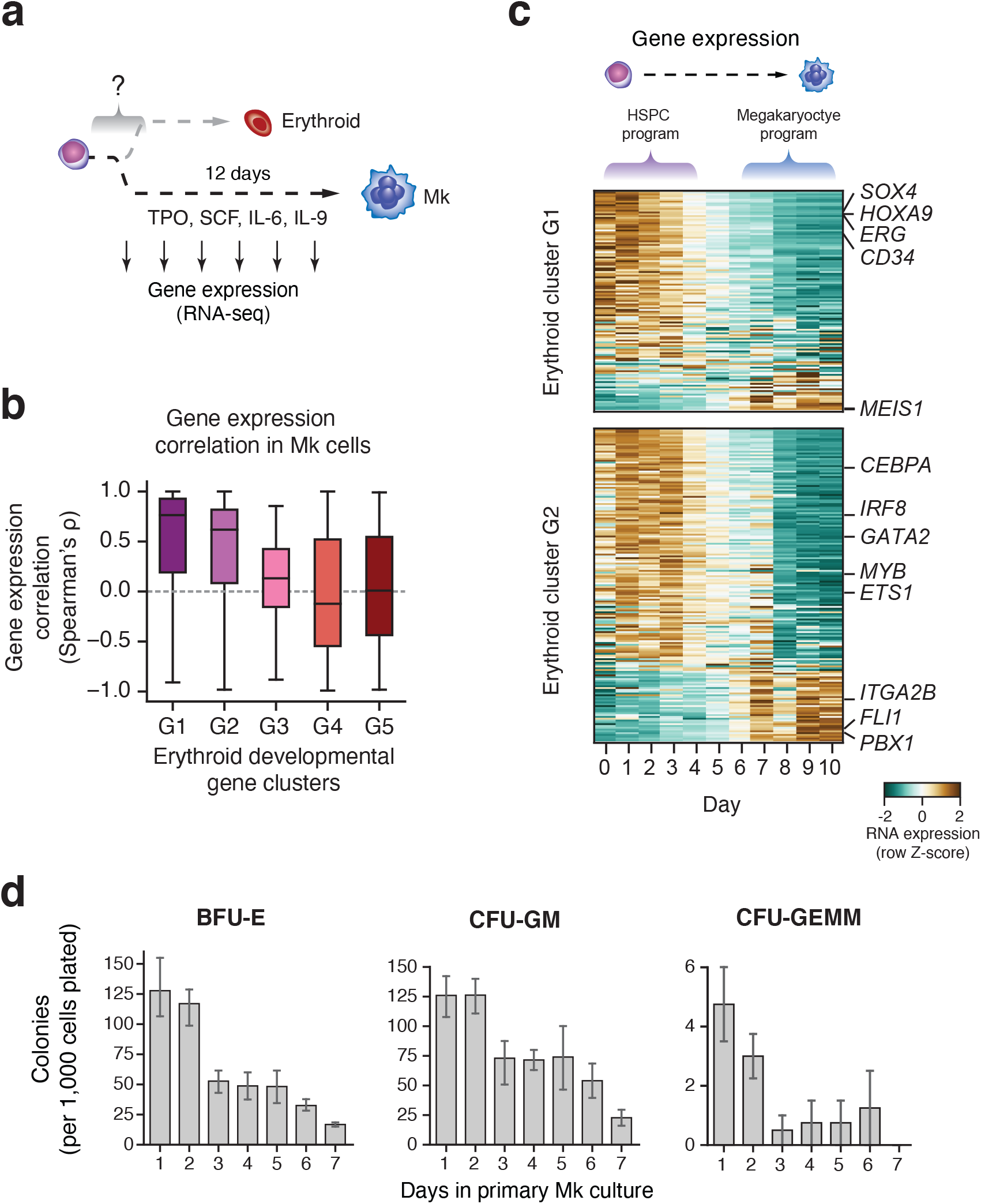
A shared transcriptional program drives the exit from HSPC state early in erythropoiesis and megakaryopoiesis. (a) Dense RNA-seq time course during *ex vivo* megakaryopoiesis induced from CD34^+^ HSPCs. (b) Correlation of gene expression profiles between erythropoiesis and megakaryopoiesis across the erythroid gene clusters G1-G5. (c) Expression profiles during megakaryocytic development, ordered by their correlation score to their erythroid counterparts from erythroid clusters G1 and G2. (d) Lineage potential assay during *ex vivo* megakaryopoiesis whereby erythroid potential was assessed by subjecting cells to a secondary erythroid culture (left). Frequency of CD235a^+^ erythroid cells after 12-day culture into secondary erythroid media (right).

Taken together these data describe the sequence of developmentally related changes in both the *cis*- and *trans*-environment as the regulatory landscape of the erythroid development traverses from a lineage-permissive program to a defined erythroid-specific signature. Expectedly, activated DHSs (clusters E4-E5) were found to preferentially harbor red blood cell-related GWAS variants (1.36-fold enrichment over all detected DHSs), highlighting their functional role in regulating erythropoiesis (**Supplementary Figure 6**).

**Figure 6.**
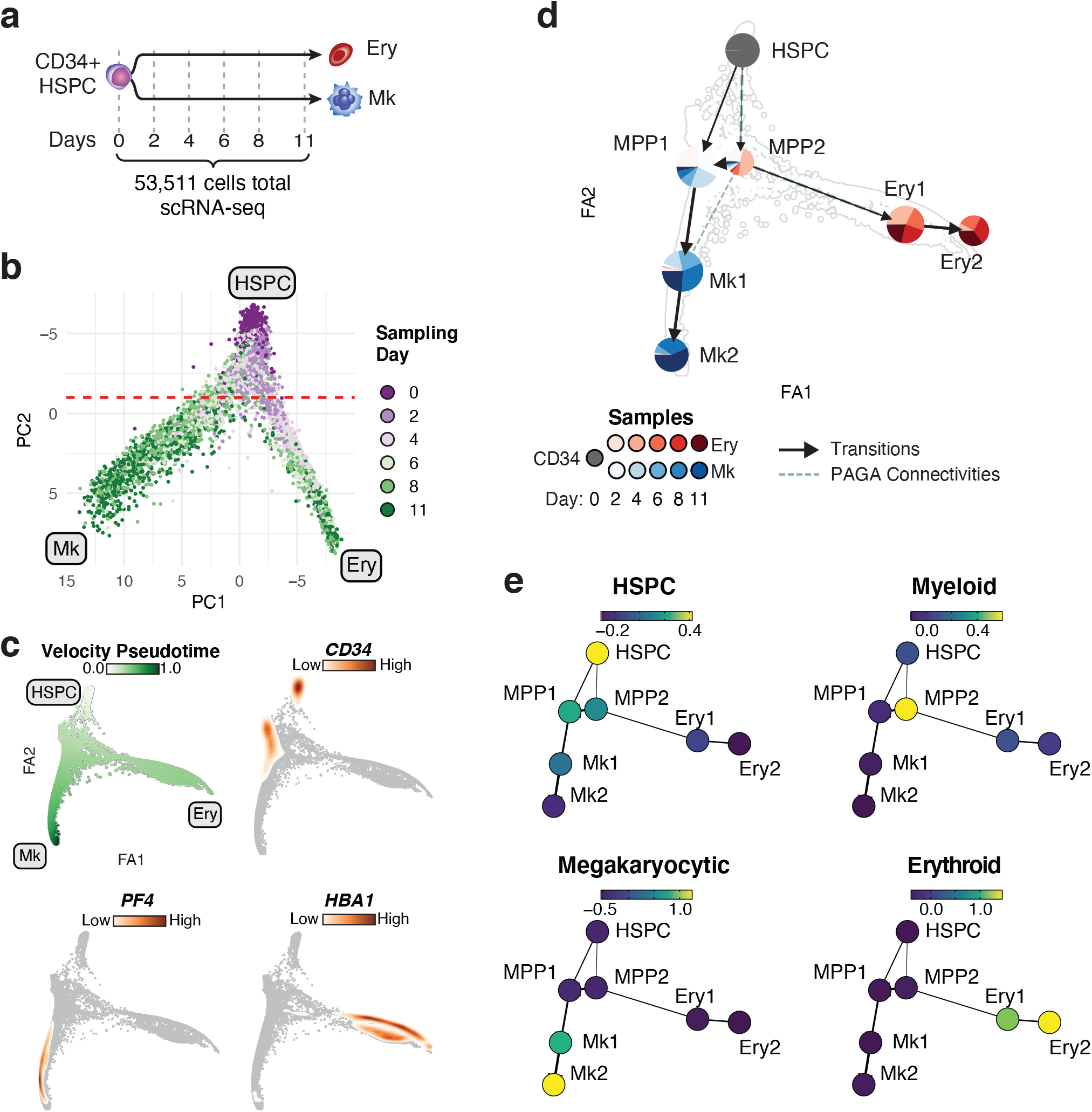
Single-cell gene expression dynamics demonstrate distinct cell states during erythroid and megakaryocytic differentiation. (a) CD34^+^ cells from a single donor were *ex vivo* differentiated towards the erythroid and the megakaryocytic lineage. Including an uncultured sample from the same donor (CD34^+^ day 0), ∼50,000 cells were totally sampled during 5 time-points from each lineage and subjected to single-cell RNA-seq. (b) PCA using the top 2000 variable genes. Cells are colored by the sampling day. Terminal erythroid (Ery) and megakaryocytic (Mk) states, as well as initial HSPC state are annotated, respectively. Red line denotes where the 90% of the cells sampled prior to day 4 are located (above line). (c) RNA velocity-based pseudotime or gene expression density on Force-Atlas graph of all cells based on RNA velocity estimates of top 2000 variable genes. (d) Cell populations from collapsed L eiden clusters on FA graph. Sample composition of each population is represented as a pie chart. Arrows denote RNA velocity derived transitions between populations. Dashed lines represent PAGA connectivities. (e) Average expression intensity of gene sets representative of HSPC, Myeloid, Erythroid and Megakaryocytic states on identified cell populations.

### Connecting individual DHSs to genes

The overall dynamics of chromatin accessibility for individual DHSs closely mirrored that of the expression of nearby genes. To formulate this, we performed an enrichment test to investigate the DHS landscape around a gene promoter. Interestingly, we found that developmentally regulated genes are enriched for DHSs with a similar developmental profile (**Figure 2c**). For example, early closing genes (cluster G1) are significantly enriched for cluster E1 DHSs. Noteworthy, transient genes of cluster G3 are harboring DHSs belonging to both late closing DHS cluster E3 and early activated erythroid DHS cluster E4, suggesting that the transient nature of these genes is a result of the combinatorial activity of a closing and an opening chromatin landscape.

Because of fine-resolution afforded by our dense sampling approach, we sought to quantify the extent of genome-wide coactivation patterns that could potentially comprise physical regulatory links between DHSs and their target genes by correlating the temporal expression patterns of a gene to nearby (±1 Mb from TSS) developmentally regulated DHSs given that the majority of transcriptional enhancers are located within this range from the target promoter^31^. This analysis identified 41,625 connections (absolute Pearson correlation coefficient *r* > 0.7), with the vast majority of gene-DHS links occurring within 50 kilobases of the transcription start site (**Figure 2d**). Overall, we connected 80.4% (4,640) of the developmentally regulated genes with ≥1 DHS and 86.8% (10,247) of changing DHSs were linked to ≥1 developmentally regulated gene. While on average 93.6 DHSs reside within ±1 Mb of a given gene, only 9 DHSs (±8 SD) were found to be linked with a changing gene. This allowed us to identify *cis*-regulatory inputs at much higher resolution than typically afforded by standard chromatin conformation-based methods^32,33^. Specifically, using previously published Hi-C data derived from *ex vivo* cultured erythroid progenitors we were able to predict chromatin loops only down to 70kb (**Supplementary Figure 7**).We therefore sought to functionally validate these associations by genetic perturbation of gene-DHS links.

**Figure 7.**
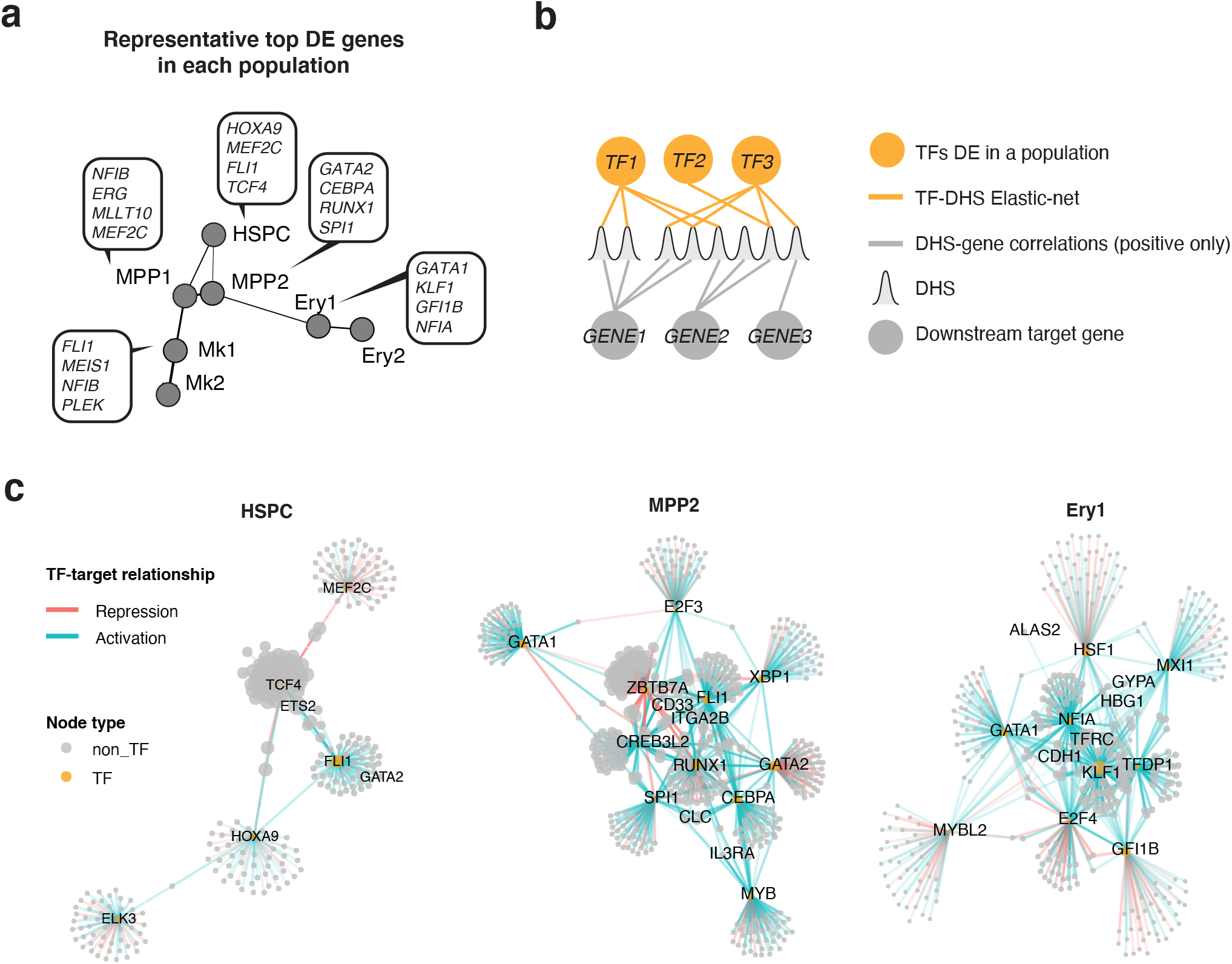
Defining of stage-specific erythroid regulatory networks. (a) Representative top differentially expressed TF genes between populations as identified by Wilcoxon sum rank test. (b) Schematic diagram of the regulatory network construction logic. Differentially expressed TFs among single-cell populations are assigned to their target genes based on elastic-net regression results. DHSs are linked to their target genes using correlation between population level DHS density and gene expression. (c) Regulatory networks initiated by TFs specific to each population. Up to top 50 target genes are shown for each TF. Transcription factors and select marker genes are annotated.

We focused on the cis-elements predicted to regulate the expression *CDH1*. CDH1 is a cell surface marker with expression restricted to erythropoiesis among the hematopoietic populations^34^ and known to be implicated in erythroid development and maturation^35,36^. Specifically, we genetically disrupted two DHSs (HS1 and HS2) highly correlated with *CDH1* expression (r=0.939 and 0.976, respectively), situated upstream (5kb and 12kb, respectively) of the promoter of *CDH1* using TALE-nucleases^37,38^ (**Figure 2e** and **Supplementary Figure 8**). Homozygous deletion of these DHSs as well as the promoter in the human derived erythroid progenitor cell line HUDEP-2 where these DHSs are also active, resulted in complete ablation of the *CDH1* expression as determined by flow-cytometry (**Figure 2f**). These results suggest that both elements as predicted by the correlation analysis as regulators of *CDH1*, indeed drive the expression of the gene and their deletion confers effects similar to the deletion of the gene promoter.

Overall, these findings suggest that the majority of changes in transcription during development are regulated by a limited number of *cis*-regulatory inputs, situated within close proximity to the genes they regulate.

### Distinct and sequential regulatory modules encode developmental stages

Clustering of dynamically changing DHSs revealed that chromatin activated at different stages of hematopoiesis display differential enrichment for transcription factor recognition sequences, indicating stage-specific transcriptional regulation of *cis*-elements. This, however, does not resolve the temporal interactions between individual DHSs and individual *trans*-regulators and how this relationship shapes the developmental response of a DHS. Given the observed global correlated changes between the transcription factor expression levels and the accessibility of the DHSs containing their cognate recognition sequences (**Figure 3a**) we sought to quantify the contribution of individual TFs to the dynamic changes in DNase I density at individual regulatory *cis*-elements. We capitalized on our dense sampling approach and applied a regression strategy where the activity of an individual regulatory element (i.e. DNase I cleavage density) is modelled as a function of the gene expression profiles of developmentally regulated TFs with a compatible recognition sequence harbored within each DHS (**Figure 3b** and **Methods**). We controlled for weak and ambiguous association of TFs recognizing degenerate motifs using elastic-net regularization (**Methods**). We applied this approach to all of the 11,805 dynamically changing DHSs, identifying 11,734 (>99% of changing DHSs) with at least one explanatory TF regulator (**Methods**) and 88 developmentally regulated TFs associated with at least one DHS, where the regression coefficients broadly correspond to the strength of association of a TF with an individual DHS (**Supplementary Figure 9**). Overall, 5 TFs on average, were positively associated with each DHS, suggesting that a small subset of TFs regulate the developmental activity of individual *cis*-elements. We then evaluated whether the regression results hold any predictive capacity against the frequency of motifs for the same TFs. Using a naïve Bayes classification model (Methods) we tried to predict the cluster each DHS belongs to by supplying either the occurrences of individual TF motifs or the elastic-net regression coefficients for each TF. We found that elastic-net regression coefficients provide a 1.77-fold accuracy over TF motif counts (62% vs. 35% accuracy rate) in predicting the DHS cluster (**Supplementary Figure 10**). This suggests that the developmental response of a DHS is shaped by co-regulated transcription factors that occupy the DHS rather than the absolute frequency of binding TFs as determined by the TF recognition sequences harbored in a DHS.

We next asked to what extent the activity of DHSs with similar temporal accessibility patterns are regulated by a coherent set of TF regulators. We selected 52 TFs positively associated with at least 200 DHSs and performed unsupervised hierarchical clustering based on their regression coefficients computed for each DHS (**Figure 3c** and **Methods**). This analysis resolved the temporal associations between transcription factors and their target DHS into a sequence of five discrete and largely non-overlapping regulatory modules, reflective of developmental stages of erythropoiesis (Figure 3d). Module 1 consists of known HSPC transcriptional regulators (e.g. ERG^28^, MEIS1^39^, MYCN^40^) which are positively associated with early closing DHSs in clusters E1 and E2. In modules 2 and 3, transcription factors associated with commitment of hematopoietic progenitors to the different myeloid lineages (e.g. CEBPA^41^, MYB^42^, FLI1^43^, RUNX1^44^) interact with DHSs in clusters E2 and E3. Modules 4 and 5 define the erythroid-specific regulatory landscape as known erythroid regulators (e.g. GATA1^45^, KLF1^20^, RXRA^46^ and FOXO3^47^) positively interact with activated DHSs in clusters E4 and E5.

Plotting the fraction of DHSs in each cluster positively associated with each TF (**Figure 3d**) highlights the major drivers of chromatin accessibility in each developmental stage. Particularly, ERG appears as a major regulator of the HPSC stage as it is positively associated with ∼25% of DHSs in clusters E1 and E2. Although ERG has been long implicated in HSPC regulation, it is only recently its role as a critical regulator of HSPC survival has been appreciated^48^. Interestingly, KLF12 also appears to share a significant proportion of the early chromatin landscape, although its role in HSPC regulation is not fully elucidated. Overexpression of the critical HSC regulator Evi-1 in mice, resulted in maintenance of the quiescent phenotype of murine HSCs along with the more than 12-fold increase in *Klf12* expression^49^. In another experiment, sustained expression of *Hlf* in mice also resulted in enrichment of *Klf12* in more primitive hematopoietic compartments^50^, thus implicating KLF12 in the HSPC regulation. Apart from the canonical erythroid transcription factors, we identified MXI1 among the top regulators of the erythroid chromatin landscape. Knockdown of *Mxi1* in mice, blocks chromatin condensation and impairs enucleation of mouse erythroblasts, highlighting the role of MXI1 in erythroid maturation^51^. Additionally, we find CTCF to be positively associated with a large portion of the erythroid-specific chromatin (∼25% of DHSs in clusters E4 and E5), while we find strong enrichment for DHS harboring CTCF motifs in predicted chromatin loops from Hi-C data generated from *ex vivo* cultured human erythroid progenitors (**Supplementary Figure 11**). These findings are consistent with the evidence highlighting the role of CTCF in establishing the erythroid-specific chromatin landscape^52,53^.

Taken together, these findings illustrate the dynamic interaction of the *cis*- and the *trans*-regulatory landscape during erythropoiesis and their organization into well-defined and discrete regulatory modules of associated DHSs with their cognate transcription factors, reflecting distinct stages of erythroid development.

### A sequence of abrupt lineage restriction events marks erythropoiesis

The organization of chromatin and transcription factors into defined regulatory modules corresponding to distinct stages of erythropoiesis indicates a functional relationship between lineage potential and module activity. To gain insight into whether these modules underpin lineage decision events we determined the lineage potential of the erythroid cultures by daily sampling a population of cells and assaying their multipotent and unipotent capacity for different myeloid lineages (**Figure 4a** and **Supplementary Figure 12a**). Total number of colonies declined with the progress of differentiation resulting in an abrupt depletion of total progenitors on day 6 of differentiation (**Supplementary Figure 12b**). After 4 days of exposure to erythroid media, the most primitive and multipotent colonies (CFU-GEMM; granulocytic, erythroid, monocytic, megakaryocytic) were no longer detected (Figure 4b). Day 6 marked a second event of restriction of the fate potential as all unilineage colonies were no longer detected in the cultures. Specifically, frequency of erythroid progenitors (BFU-E) rapidly declined from day 5 to day 6 (**Figure 4c**). Similarly, granulocytic/monocytic progenitors (CFU-GM) were depleted by day 6 of erythroid differentiation (**Supplementary Figure 12c**). Notably, none of the changes in clonogenic capacity were associated with any changes in the growth rate of the parental erythroid cultures, which remained constant throughout the differentiation (**Supplementary Figure 12d**), suggestive of an independent mechanism regulating this shift in progenitor population.

In addition to the above lineages, we specifically tested for the ability to differentiate into megakaryocytes during erythroid development by transferring cells, on a daily basis, from the primary erythroid culture to megakaryopoiesis-inducing suspension cultures and tested for their ability to give rise to CD41^+^ megakaryocytic populations (**Supplementary Figure 13a** and **Methods**). Consistent with the overall lineage restriction observed during colony-forming assays, erythroid cultures completely lose megakaryocytic potential on day 6 of the differentiation (**Figure 4c** and **Supplementary Figure 13b**).

The rapid changes observed in clonogenic capacity correspond to the transitions in the activity of regulatory modules (**Figure 4d**). Early depletion of primitive multipotent CFU-GEMM progenitors is concomitant with the transition from the HSPC-related modules (modules 1 and 2), while the decline of unipotent progenitors of all detectable myeloid lineages (granulocytic/monocytic, erythroid, megakaryocytic) coincides with the transition from a program with a broader myeloid signature to erythroid specific *cis*- and *trans*-landscape. Furthermore, because these rapid lineage restriction events are not associated with other abrupt changes in morphology or cell growth (**Supplementary Figure 13d**), these data suggest that the mechanism responsible for the exit from the progenitor stage is decoupled from maturation progress.

### Exit from the HSPC-related transcriptional program is shared between erythropoiesis and megakaryopoiesis

The silencing of the HPSC regulatory modules prior to lineage commitment suggested that exit from the progenitor state is necessary for erythroid commitment to proceed. We therefore asked whether this represents a canonical feature of hematopoietic development to any lineage. To investigate this, we focused on megakaryocytic differentiation, a process that shares both close common developmental origins^54^ and key TF regulators with erythropoiesis^55^.

We induced *ex vivo* megakaryocytic differentiation and performed dense sampling of gene expression during development (**Figure 5a** and **Methods**). Developmentally regulated genes during megakaryopoiesis exhibit largely bipartite profiles similar to those observed during erythropoiesis (**Supplementary Figure 14**). To determine whether the transcriptional changes associated with exit from HSPC state during erythropoiesis are shared with megakaryopoiesis we examined the expression profiles of erythroid developmentally regulated genes during megakaryocytic differentiation. We observed highly correlated global expression profiles for early silenced transcripts (erythroid clusters G1 and G2) between the two lineages (median Spearman’s *ρ*=0.76 and 0.62, respectively) (**Figure 5b**), with the exception of key regulators and canonical markers of megakaryopoiesis (*MEIS1, FLI1, PBX1, ITGA2B*, etc.) (**Figure 5c**). In contrast, correlation for erythroid clusters G3-G5 was low (median Spearman’s *ρ ≤ 0*.*13)*.

Similar to erythropoiesis, we found that early downregulation of HSPC-related gene signature is associated with abrupt restriction of alternate lineage potential during megakaryopoiesis. Specifically, we found that cells sampled beyond day 3 of differentiation exhibit a reduction in both multipotent and unipotent progenitors of the erythroid and granulocytic/monocytic lineage (**Figure 5c** and **Supplementary Figure 15**). This observation is in line with the fact that megakaryopoiesis does not exhibit transient activation of myeloid gene program as exit from HSPC is rapidly succeeded by a megakaryocyte-specific gene signature. This finding is compatible with the recently revised hematopoietic tree according to which megakaryocytes directly emerge from the primitive HSPC compartments bypassing the common myeloid progenitor^56–58^.

Conclusively, these results indicate the existence of a shared mechanism between erythropoiesis and megakaryopoiesis driven by a common set of TFs which mediates the exit from HSPC state signaling differential lineage potential response for erythropoiesis and megakaryopoiesis.

### Transient acquisition of a myeloid signature precedes erythroid commitment but not megakaryocytic

Genomic and functional findings on population-level during *ex vivo* erythropoiesis suggest that erythroid development transitions through a state with permissive alternate lineage potential prior to erythroid commitment whereas megakaryocytes appear to rapidly commit after exit from HSPC. As lineage decision events resolve in individual progenitors, we sought to resolve the fate commitment kinetics and the differentiation trajectories of erythropoiesis and megakaryopoiesis by jointly analyzing transcriptional dynamics in single cells along the two lineages. To this end, we analyzed transcriptional changes from more than 50,000 single cells sampled from frequent intervals along both the *ex vivo* erythroid and megakaryocytic differentiation (**Figure 6a**). Overall, we found that single-cell gene expression profiles to be highly concordant with total RNAseq data as aggregated gene expression from single-cell RNA-seq correlated very well with RNA-seq performed in bulk cells (**Supplementary Figure 16**).

Principal component analysis (PCA) using the top 2,000 variable genes readily resolved the two primary axes of differentiation. Trajectories from HSPC to terminally committed lineages are resolved along PC2 while PC1 distinguishes the erythroid and megakaryocytic terminal fates (**Figure 6b**). Furthermore, PCA highlights the lineage commitment timepoint as cells sampled on day 4 and thereon, from either culture, already exhibit distinct topologies on the PCA projection. In order to infer rate of transcription and derive the direction of differentiation, we capitalized on splicing kinetics (RNA velocity) to derive latent pseudotime (**Figure 6c**). Overall, pseudotime correlated well with actual sampling time (Pearson’s *r*=0.725).

Cells were clustered using Leiden community detection algorithm^59^ and based on their between affinities cell clusters were collapsed to 7 distinct populations corresponding to discrete developmental stages. Developmental pseudotime and transitions between populations were inferred based on RNA velocity (**Figure 6d** and **Supplemental Figure 18a-c**). Overall, we found the populations to be highly homogeneous in terms of *ex vivo* culture sample composition. Not unexpectedly, we observed higher sample admixture in populations corresponding to eary time-points, consistent with the notion that cells at this stage had yet to establish lineage fate (**Supplementary Figure 18d**).

Using lineage trajectories inferred from transcriptional transitions between populations, we identified two major pathways starting from a cluster with HSPC signature (HSPC cluster) and leading to terminal megakaryocytic and erythroid fates (**Figure 6c**). Transitions from HSPC cluster to lineage specific clusters involves two clusters with progenitor gene signature (MPP1, and MPP2) each of them stemming from the HSPC cluster. MPP1 consists primarily of cells sampled from megakaryocytic cultures while ∼25% of the cells in the population are derived from day 2 of the erythroid differentiation. MPP1 maintains a broader early progenitor signature (**Supplementary Figure 18e, f**) and transitions to a population with early Mk signature (Mk1) which eventually gives rise to mature megakaryocytic population (Mk2). MPP2 is composed almost exclusively of early (day 2 and day 4) erythroid cells and appears as a nodal cluster with affinities to both the early erythroid cluster as well as the Mk primed MPP1. Importantly, MPP2 exhibits gene expression signature characteristic of various myeloid subtypes (**Supplemental Figure 18e-f**) expressing critical myeloid regulators alongside megakaryocytic and erythroid ones (**Supplementary Table 6**). Stage-specific TF network reconstruction using TF-DHS and DHS-gene assignments (**Figure 7a,b**), reveals the myeloid signature is orchestrated by a core network of critical myeloid regulators (SPI1/PU.1, CEBPA, and FLI1), specific to MPP2 (**Figure 7c**). This is compatible to single cell TF protein dynamics during *ex vivo* erythropoiesis, demonstrating that multiple TFs of alternate hematopoietic lineages are active in early progenitors prior to emergence of CFU-e populations^60^. Furthermore, early erythroid progenitors display affinities with myeloid and basophilic lineages transiently emerging in the culture. Here, in an attempt to identify the origin of this myeloid population present in our experiments, we performed FACS timecourse for the myeloid marker CD33 during both erythropoiesis and megakaryopoiesis. We find that the CD33^+^ population a subset of CD34^+^ HSPCs as >80% of uncultured CD34^+^ are also positive for CD33. During erythroid differentiation, we find that cells transiently undergo a state of CD33^+^/CD117^+^, whereby day 6 the majority (∼70%) has transitioned to CD33^-^/CD117^+^ (**Supplementary Figure 19**). In contrast, expression patterns of CD33 and CD41 during *ex vivo* megakaryopoiesis are mutually exclusive, confirming the erythroid-specific origin of the myeloid population.

In order to compare our findings to steady state hematopoiesis, we analyzed previously published single-cell RNAseq data from FACS fractionated BM-derived hematopoietic populations^61^. Trajectory inference on Force Atlas embedding using PAGA and DPT pseudotime analysis revealed two major differentiation pathways originating from a developmentally primitive population with HSPC. One with defined erythroid signature, and one exhibiting a myeloid gene expression profile (**Supplementary Figure 20a**). Although we were able to detect a few cells with megakaryocytic signature concentrated close to HSPCs, their population is very small and no distinction between primitive and mature megakaryocytes could be detected. Additionally, gene expression patterns of megakaryocytic markers and TFs are not well defined to infer differentiation trajectory (**Supplementary Figure 20a**). Upon determination of cell clusters (**Supplementary Figure 20b**) we detected a population of cells which expresses several myeloid markers and particularly those of basophils (e.g., *LMO4, CLC*), and appears to originate from two populations early on the erythroid trajectory (**Supplementary Figure 20b**). In order to compare gene expression profiles along the erythroid trajectory inferred from either *ex vivo* differentiated erythroid cells or BM fractionated populations we correlated gene expression profiles from 1,000 top highly expressed genes in both datasets. This revealed that *ex vivo* erythropoiesis recapitulates exceptionally well the gene expression dynamics from native erythroid populations with median Spearman’s *ρ*=0.81 (**Supplementary Figure 21**). These results further support our population-level findings about the transient emergence of a population during erythroid development that maintains myeloid capacity. The affinity between the basophilic lineage and the erythroid has been previously described^61,62^ and it has been suggested that basophils derive from erythro-myeloid progenitors^63^.

## Discussion

Here, we systematically link individual transcription factors and their target *cis*-elements along *ex vivo* human erythropoiesis, resolving how these elements organize temporally, encoding lineage commitment and differentiation during hematopoiesis. More recently, multiple efforts have extensively studied the individual (*cis*- and *trans*-) regulatory components involved in erythropoiesis^22^ as well as other diverse hematopoietic lineages^9,64–67^. The bulk of these efforts however base their findings either on immunophenotypically defined hematopoietic populations, or single-cell dissection of steady state heterogeneous sources, where developmental relationships between cells within a heterogeneous steady-state population can only be inferred^13,15,61,68^. In this work we overcome the limitations associated with immunophenotypic isolation of hematopoietic populations^57,69^ which often fail to capture transient or rare populations, while enrichment for specific populations is entirely dependent on the immunophenotypic panel used for fractionation^70^. By capitalizing on the continuity of the differentiating populations during *ex vivo* erythropoiesis we finely map chromatin accessibility and gene expression dynamics enabling the direct and repeated measurement of the dynamic epigenetic landscape along a defined lineage trajectory. In addition, a dense sampling approach enables the unbiased detection of transient events occurring over short intervals that would otherwise be missed by sparse sampling methodologies.

Through integrative analysis of chromatin accessibility and gene expression during erythropoiesis we draw thousands of links between individual distal regulatory elements and their target genes at much higher resolution than that afforded by other methods. This approach revealed a sequence of discrete, non-overlapping regulatory modules comprising of interacting transcription factors and individual *cis*-regulatory elements, corresponding to distinct stages of erythroid development. Strikingly, the transition between the activity of these modules coincides with a sequence of experimentally validated rapid lineage restriction events. We found that the exit from the program associated with the HSPC state occurred independent of lineage outcome, as it was also identified during *ex vivo* megakaryopoiesis. Moreover, comparison of developmental transcriptomics of single cells along erythropoiesis and megakaryopoiesis reveals that exit from HSPC occurs over the same developmental interval for both lineages, indicative of a mechanism independent of the cytokine environment. This finding adds to previous reports that activation of murine bone marrow HSCs with different lineage cytokines induces a common repression mechanism of HSC signature while activates genes implicated in differentiation in a cytokine independent fashion^71^

Upon exit from the HSPC state we found the two lineages to exhibit differential commitment kinetics. Erythroid differentiation maintains a broader myeloid lineage capacity (Ery, G/M, Mk) prior to erythroid commitment as a result of a transient upregulation of a regulatory program involving canonical myeloid transcription factors (FLI1, SPI1, C/EBPs, GATA2, etc.). Network analysis in the progenitor stage prior to erythroid commitment, demonstrates that FLI1 is a central TF with extensive affinities to other transcriptional regulators, ultimately gatekeeping the fate choice between the megakaryocytic and erythroid lineage. There are several lines of evidence from single-cell assays in both mouse and human hematopoiesis suggesting that erythroid, megakaryocytic and basophilic lineages emerge from a shared population^13,16,60,64,72,73^, while mass cytometry dynamics of lineage-specific transcription factors ascribe FLI1 the role of “gatekeeper” between the erythroid and the megakaryocytic fate^60^. The findings presented here, however, demonstrate that this lineage-permissive transcriptional program is restricted only to erythropoiesis. This is compatible with previous experimental evidence demonstrating the affinity of basophilic lineage to the erythroid branch^61^, specifically. Additionally, results from transgenic mice lacking a set of the C/EBP family of myeloid regulators that exhibit decreased erythroid output^74^. In contrast, transcriptional, functional and phenotypic evidence from *ex vivo* megakaryopoiesis presented here, suggests rapid megakaryocytic commitment upon HSPC exit. These results align well with the growing evidence suggesting that megakaryocytic commitment is occurring earlier compared to erythroid fate^75^ and that megakaryocytic lineage arises directly from the primitive hematopoietic compartments^57,76–78^.

In order to reconcile our findings on lineage commitment from bulk populations with transcriptional dynamics from individual cells we compiled one of the most comprehensive analyses of single-cell gene expression along a closely monitored developmental system, so far. Furthermore, this dataset represents the first, to our knowledge, single-cell description of gene expression dynamics along the *ex vivo* megakaryocytic development from purified HSPCs. Although different approaches have been suggested to enrich for megakaryocyte and platelet biased progenitors and dissection of the bipotent MEPs^79–81^ there is no consensus purification scheme to isolate cells at different stages of megakaryocytic development with the resolution available for erythroid development^82,83^. This is primarily due to the rarity of Mk cells in the bone marrow^84,85^ and the fragility of the mature large endomitotic megakaryocytes. Here, however, we present an unbiased global view of gene expression and lineage commitment dynamics of megakaryocytic development based on equiproportional sampling of populations along megakaryopoiesis.

Overlaying the information of sampling timepoint of each population allowed us to match shifts in TF expression in single-cells to the regulatory programs identified from our population-level experiments. Strikingly, our single-cell based observations recapitulate both our *ex vivo* population-based findings as well as single-cell transcriptional dynamics from bone marrow fractionated populations with remarkable fidelity. This suggests that highly synchronized rapid shifts in gene expression levels of lineage regulators across individual cells, occurring over short intervals of developmental time, underpin the changes observed in bulk populations. This contrasts the current sentiment hinging on observations from single-cell analyses where variability in the chromatin and transcriptional landscapes among steady-are interpreted as gradients of continuous regulatory states^13,14,68,86^.

Here we present novel insights into the developmental regulatory dynamics during hematopoiesis illuminating mechanisms of lineage commitment, unable to interrogate previously due to sampling biases and limitations. Although we draw parallels with steady-state *in vivo* derived data, the artificial nature of the *ex vivo* culture systems can be a confounding factor. Nevertheless, we highlight the utility of *ex vivo* development systems in studying rare or otherwise inaccessible populations *in vivo* and provide a generalizable framework of how interactions between the *trans*-environment and the chromatin instruct fate choice and lineage commitment during development. Additionally, the dense sampling and the systematic linkage between distal elements and target promoters results in high-resolution maps charting the stage-specific activity of regulatory elements. Such elements can prove particularly useful in transgene-based therapies where the efficacy of these methods relies on the precise modulation of gene expression in a developmental and lineage-specific manner.

## Author contributions

G.G., M.I., and N.P. performed the experiments. G.G., A.N., T.S., and J.V. performed data analysis. G.G., N.P., and J.V. designed the experiments. J.S., M.Y., and J.V. supervised the study and provided consultation. G.G., and J.V. wrote the manuscript. J.V. conceived the study. All authors approved the manuscript.

## Acknowledgments

The authors would like to thank Daniel Bates, Morgan Diegel, Douglas Dunn, Fidencio Neri, Ericka Otterman, Shinny Vong, and Alister Funnell from the Altius Institute for Biomedical Sciences for help in sequencing and genome editing. We also thank John Lazar and Thalia Papayannopoulou for their critical review on the manuscript.

## Data availability

All DNase I and RNA-seq data produced as part of this manuscript is freely available by request and has been submitted to SRA. Adult erythroid Hi-C data was downloaded from ERA (accessions SRX3058042 and SRX3058043). Adult CD43+ HSPC Hi-C data obtained from ENA (accession ERR436024). Human bone marrow single-cell data from Pellin *et al*., *2019, Nat. Comms*. was download from ERA (accessions SRX4455330, SRX4455331, SRX4455332, SRX4455333, SRX4455334, SRX4455335 and SRX4455336).

## Code availability

All scripts and code used to process and analyze data herein are available upon request.

