## Supplemental methods and figure legends for "Chromatin dynamics during hematopoiesis reveal discrete regulatory modules instructing differentiation"

### Supplementary material for: "Chromatin dynamics during hematopoiesis reveal discrete regulatory modules instructing differentiation"

Grigorios Georgolopoulos, Nikoletta Psatha, Mineo Iwata, Andrew Nishida, Tannishtha Som, Minas Yiangou, John A. Stamatoyannopoulos, Jeff Vierstra

#### Materials and Methods

***Ex vivo* erythropoiesis.** For the *ex vivo* induction of erythroid and megakaryocytic differentiation, Human CD34+ enriched PBMC (>90% purity) from 3 different GCSF-mobilized healthy adult donors (Fred Hutch Cancer Research Center – Cooperative Centers of Excellence in Hematology Core B) were used. Prior to culture, cells were thawed rapidly in a 37ºC water bath and cultured overnight in recovery media containing IMDM, supplemented with StemSpanTM CC110 (Stemcell Technologies). For erythroid differentiation an established 3-step differentiation scheme was used[1]. Briefly, CD34+ cells were cultured for 7 days in IMDM media containing 0.1 µg/mL rhSCF (PeproTech), 0.005 µg/mL rhIL-3 (PeproTech), 3 U/mL rhEPO (PeproTech), 5% human AB plasma, 2 U/mL heparin (PRODUCT INFO), 0.01 mg/mL Insulin, 0.33mg/mL holo-transferrin (Millipore-Sigma), 1µM hydrocortisone (Millipore-Sigma), 1x Penicillin/Streptomycin (ThermoFisher Scientific), followed by culturing in the same media for 4 more days without IL-3 and hydrocortisone for 4 days.

***Ex vivo* megakaryopoiesis.** CD34+ HSPCs were differentiated to the megakaryocyte lineage by culturing for 11 days in IMDM based media containing 30ng/mL rhTPO (PeproTech), 1ng/mL rhSCF (PeproTech), 7.5ng/mL rhIL-6 (PeproTech), 13.5ng/mL rhIL-9 (PeproTech), 20% BIT (StemCell Technologies), 40µg/mL LDL (Millipore-Sigma), 0.05mM beta mercaptoethanol (Millipore-Sigma), and 1x Penicillin/Streptomycin (ThermoFisher Scientific).

**HUDEP-2 Culture.** HUDEP-2 cell culture HUDEP-2 cells were maintained in StemSpan H3000 medium (Stem Cell Technologies) supplemented with 100 ng/ml hSCF (Peprotech), 3 IU/ml erythropoietin (Peprotech), $10^{-6}$ M dexamethasone (Millipore Sigma) and 1 mg/ml doxycycline (Millipore Sigma).

**Genetic knock-outs in HUDEP-2 cells.** All TALEN pairs were assembled according to established protocols[2,3]. Sequences of the TALEN binding domains are listed in Supplementary Table 7. In vitro transcription of TALEN-mRNA was performed by the T7 mScript™ Standard mRNA Production System (CELLSCRIPT, C-MSC100625), including 5'-capping and poly-A addition reactions. Electroporation was performed as previously described[4]. Briefly, 2µg of each TALEN monomer mRNA were transfected in HUDEP-2 cells using a BTX electroporator (ECM 830, Holliston, MA) in 100 µL of BTX Express electroporation solution. Double knock out clones were detected post electroporation and single cell sorting (MoFlo Astrios, Beckman Coulter) by Out-Out (CDH1 promoter and HS1) or In-Out PCR (HS2). PCR Primer sequences are provided in Supplementary Table 7.

**Colony forming assays.** Colony forming assays were performed by sampling ~1000 cells daily from either erythroid or megakaryocytic primary cultures from 2 donors and plating each in duplicates in 35mm dishes containing MethoCult H4435 (StemCell Technologies). After two weeks, colonies were identified and scored under a dissection microscope.

**Suspension lineage potential assays.** For megakaryocytic potential assays, ~100,000 cells were sampled daily from erythroid cultures until day 7. After removing primary media, cells were transferred to secondary suspension cultures containing megakaryocytic media as described above and allowed to grow for two weeks. Megakaryocytic potential was estimated by the frequency of $CD41^+$ cells with flow cytometry on day 12 of the secondary megakaryocytic culture. Similarly, for erythroid potential assays during megakaryocytic differentiation, ~100,000 cells were sampled daily until day 7 from primary megakaryocytic differentiation and transferred to secondary erythroid media. After two weeks in the secondary media, erythroid potential was assessed by the frequency of $CD235a^+$ cells present with flow cytometry.

**Flow cytometry.** Approximately 100,000 cells each day per culture replicate were harvested. Media were removed by washing with staining buffer (PBS + 0.25% BSA) and cells were labelled according to product specifications, incubating at 4ºC. Cells from erythroid differentiation were stained daily for CD117 PE (Clone 104D2, BD Biosciences), and CD235a (Clone GA-R2/HIR2, BD Biosciences). Megakaryocytic differentiation was monitored by staining with CD41 PE (Clone HIP8, BD Biosciences), and CD42b APC (Clone HIP1, BD Biosciences). Megakaryocytic potential of primary erythroid cultures was determined by measuring CD41a PE (Clone HIP8, BD Biosciences) expression in the secondary cultures. Myeloid populations were identified by CD33 expression staining with Alexa Fluor 700 anti-human CD33 (Clone WM53, BD Biosciences). For detection of genetic knock-out of CDH1 in HUDEP-2, APC anti-human CD324 (E-Cadherin) (Clone 67A4, Biolegend) was used. Samples were acquired on CytoFlex S (Beckman Coulter) and analyzed using FlowJo (Becton Dickinson).

**DNase I accessibility.** Erythroid cultures from 3 donors were sampled on a daily basis from day 0 to day 12 and >100,000 cells were harvested per DNase I reaction adapting the protocol from John *et al.*[5]. After removing media by washing with ice-cold PBS, cells were washed with ice-cold Buffer A (15 mM Tris-HCl pH 8.0, 15 mM NaCl, 60 mM KCl, 1 mM EDTA pH 8.0, 0.5 mM EGTA pH 8.0, 0.5 mM spermidine). Cells were resuspended in ice-cold Buffer A and equal volume of lysis buffer (0.04% de-ionized IGEPAL CA-630 in Buffer A) was added and cells were incubated at 4 ºC for 10 min. Nuclei were then moved to 37 ºC and replicates of each sample were incubated with a gradient of DNase I solution (40 IU to 100 IU of DNase I) of equal volume and the reaction was allowed to proceed for 3 min at 37 ºC. DNase I digestion was quenched by adding a volume of 5X Stop buffer (50 mM Tris-HCl pH 8.0, 100 mM NaCl, 0.1% SDS, 100 mM EDTA pH 8.0) equal to the volume of cell suspension and the reactions were incubated at 37 ºC for 60min. After incubation, 1µL of Proteinase K (Sigma) was added and the reactions were incubated at 50 ºC for 60min. Digested genomic DNA was visualized on 1.2% agarose gen and the fragment size profile was generated using the Fragment Analyzer (Advanced Analytical).

Prior to genomic library generation, fragments were subjected to size fractionation. Large DNA fragments were removed mixing the DNase I digested sample with a Polyethylene-Glycol (PEG 8000, Sigma) solution containing 8.3 mg of carboxylate-modified magnetic particles (Sera-mag beads, Thermo) to a final PEG concentration of 6% w/v. Sample was incubated in room temperature with constant mixing. Bead-bound fragments were removed by magnetic separation and a PEG solution (39.5% w/v final) containing 7.15 mg magnetic beads was added to the supernatant. After >90 min incubation at room temperature with constant mixing, the supernatant is removed by magnetic separation and discarded while bound DNA fragments are eluted from the beads. The eluate is then subjected to a second binding by mixing with a PEG solution (38.5% w/v final) containing 7.15 mg of magnetic beads. The solution is incubated at room temperature for >90 min with constant mixing. The supernatant is removed by magnetic separation and the fragments are eluted from the beads. Fragment size distribution and concentration of the fractionated sample was measured with Fragment Analyzer (Advanced Analytical). Illumina compatible, double-stranded DNA library libraries from the size fractionated samples were constructed using the ThruPLEX DNA-seq Kit (Takara Bio) according to manufacturer's instructions. DNase I-seq libraries were sequenced on NextSeq 500 (Illumina) with a 2x36bp read length. Adapter trimmed FASTQ files were aligned against GRCh38 using the BWA aligner[6]. All downstream DNase I-seq analyses were performed on DNase I hotspots (genomic regions with statistically significant enrichment in DNase I cleavage)[7,8]. Hotspots were detected using hotspot2 program (https://github.com/Altius/hotspot2).

**Total RNA sequencing.** For gene expression analysis, total RNA was collected using the mirVana RNA isolation kit (ThermoFisher Scientific) or RNeasy Mini Kit (Qiagen) from sorted (>20,000 cells) and bulk cultures (>1,000,000 cells). Illumina libraries were constructed using the TruSeq Stranded Total RNA with Ribo-Zero Globin (Illumina). Finally, libraries were quantified using Fragment Analyzer (Advanced Analytical). RNA-seq libraries were sequenced with HiSeq 4000 (Illumina) using a 2x76bp read length and alignment was performed using STAR Aligner[9] against the GRCh38 reference genome. Gene counts were obtained using featureCounts[10] and FPKM per gene were calculated using Cuffllinks[11]. Values were normalized using quantile normalization.

**Identification of developmentally regulated DHSs and genes:** To identify developmental responses in chromatin accessibility and gene expression, we employed a regression analysis strategy. A robust linear regression was performed on the quantile-normalized, mean-centered DHS density values or gene counts from 3 donors by fitting a cubic spline with 3 degrees of freedom using the lmrob function from the robustbase R package and fitted values were recorded. Statistical significance was estimated by performing a likelihood ratio test against a null model where developmental time was removed as a term. Likelihood ratio test (LTR) *p*-values were adjusted for false discovery rate (FDR) using the Benjamini-Hochberg method[12]. A refined list of the statistically significant changing DHS was obtained by setting the maximum normalized DHS density to 30 counts and $\log_2$-fold difference between the minimum and maximum daily average values to 1. Significantly changing genes were further filtered by excluding genes with maximum FPKM below 2 and those with $\log_2$-fold difference between the minimum and maximum daily average gene counts below 1.

**Identification of TADs and interaction loops from Hi-C data.** Capture Hi-C data from primary $CD34^+$ hematopoietic progenitor cells were obtained from Misfud *et al.* 2015[13]. *Ex vivo* differentiated day 11 erythroid progenitors derived Hi-C data were obtained from Huang *et* al., 2017[14]. Reads were processed with pairtools (https://github.com/open2c/pairtools) removing reads with MAPQ < 30. Contact matrices were generated with Cooler[15]. Topologically associated domains (TADs) were computed with HiCExplorer[16] on KR balanced matrices at 10kb resolution. Chromatin interaction loops were predicted using Mustache[17] from the KR balanced matrices at 10kb resolution.

**Transcription factor recognition sequence enrichments.** A list of transcription factor motif position weight matrices (PWM) was compiled from JASPAR 2018[18] and HT-SELEX derived human transcription factor binding specificity models[19]. Motifs were then scanned across the GRCh38 reference genome using FIMO[20] and motifs with nominal *p*-value $< 10^{-4}$ were mapped to DHS using BEDOPS[21]. Over- or under-representation of transcription factor motif clusters in each of the K-means DHS clusters was tested by performing a one-tailed hypergeometric enrichment test. *P*-values were adjusted using the Benjamini-Hochberg method[12].

**Elastic net regression and regulatory module identification.** To identify the transcription factors that modulate the accessibility of DHS a two-step regression with elastic-net regularization was utilized where the per-day averaged accessibility of each DHS over time was expressed as a function of the per-day averaged expression of the transcription factors with motifs found in the DHS. We considered 214 transcription factors which had maximum expression >2 FPKM and for which there was available binding motif information. An elastic net support vector machine was initially trained on a subset of the data by omitting 4 timepoints using the glmnet R package. Optimization for the mixing parameter $\alpha$ ($0 < \alpha < 1$) and the penalty stringency parameter $\lambda$ was performed with a 100-fold cross-validation of each seat of $\alpha$ (ranging from 0 to 1 with 0.01 increments) and $\lambda$ (100 equiproportional values ranging from $10^{-5}$ to $10^{5}$) parameters using the cv.glmnet function. Mean squared error (MSE) and standard error (SE) for each parameter pair tested and each DHS were recorded. Given that each DHS cluster is characterized by specific features (i.e. DHS density profile, sequence composition, and DHS overlap with other tissues) we sought to optimize elastic net parameters for each cluster, rather than overfitting each DHS individually. Therefore, for each cluster of DHS, the pair of $\alpha$ and $\lambda$ parameters which minimized the total MSE and SE for that cluster was selected and subsequently supplied into a final elastic net regression which was applied to each DHS in a cluster (E1-E5). The capacity of elastic-net TF coefficients to accurately classify the DHS into their respective *K*-means cluster (E1-E5) against the TF motif counts was evaluated using a naïve Bayes classifier using fastNaiveBayes (https://github.com/mskogholt/fastNaiveBayes). Discrete motif counts per DHS for TFs with at least one non-zero elastic-net coefficient (192 out of 214) were modelled using a Poisson fit while elastic-net coefficients for the same set of TFs were modelled with Gaussian fit and prediction accuracy was calculated as the percent of correct classifications. Transcription factors with >200 DHS positively associated (52 out of 214 TFs) were taken into consideration and a hierarchical clustering on the cosine distances of the regression coefficients was performed. Cutting the tree at the 5 highest order clades using *k*=5 resulted in 5 clusters of transcription factors which together with their associated DHS constitute a regulatory module.

**Single-cell RNA sequencing and data processing.** Erythroid and megakaryocytic cultures were induced from the same donor as described above and on days 2, 4, 6, 8, and 11 cells were harvested and stored in liquid nitrogen using CryoStor CS10 (StemCell Technologies) until library preparation. On the day of library preparation, frozen cultured cells as well as an uncultured vial of $CD34^+$ cells from the same donor were simultaneously processed. Cells were prepared for library preparation according to manufacturer's instructions using the Chromium Single Cell 3′ Reagent Kit, version 1 (10X Genomics) for $CD34^+$ and erythroid samples, except day 2. Chromium Single Cell 3′ Reagent Kit, version 2 (10X Genomics) was used for all megakaryocytic differentiation time points, and erythroid day 2. All libraries were sequenced on HiSeq 4000 (Illumina). Raw sequencing data were processed using Cell Ranger analysis pipeline v2.1.1. Reads were aligned against the GRCh38 reference genome.

**Single-cell RNA sequencing analysis of samples from primary erythroid and megakaryocytic differentiation.** RNA velocity was computed using the CLI component of velocyto[22] and Loom files with normalized transcript counts for each sample were generated with the 'run10x' command. Trajectory and pseudotime analysis was performed using the scVelo[23] and SCANPY[24]. Gene count matrix was filtered to include the top 2,000 variable genes. Moments of spliced and unspliced kinetics were calculated using the first 30 components and 30 nearest neighbors. Cells were clustered using Leiden community detection algorithm[25] and were further collapsed into biologically relevant populations. Cell connectivities, transitions, and pseudotime were computed based on RNA velocity. Developmental trajectories were inferred using PAGA[26] on velocity connectivities and transitions projected on a Force-Atlas (FA) graph embedding using a velocity pseudotime prior. Marker genes per population were identified using Wilcoxon-rank test for every population versus the immediately connected populations. Significance was called on $FDR < 10^{-5}$ and absolute $\log_2$ fold-change $\geq 1$. Overrepresentation of gene sets characteristic of specific cell-types, present within the marker genes from each population, was performed using ENRICHR[27] available at (https://maayanlab.cloud/Enrichr/) using annotated gene-sets from the Human Gene Atlas database.

**Reconstruction of stage-specific transcription factor networks.** For every differentially expressed transcription factor gene in each population, linked DHSs were called based on the elastic-net results. Subsequently, downstream target genes were identified based on positively correlated (Pearson's $r \geq 0.7$) DHS and genes. Networks were constructed using the igraph package (https://igraph.org/r/). Edge weights were calculated by standardizing the product of the elastic net coefficient and the pearson correlation value. For each node the centrality value was calculated. Network plots were generated using tidygraph (https://github.com/thomasp85/tidygraph) and ggnet2 (https://briatte.github.io/ggnet/) using the "fr" layout option.

**Single-cell analysis of human bone marrow derived data.** Raw transcript count matrix obtained from Pellin *et al.*, 2019[28] was filtered to include genes above 800 total counts and normalized to read depth and highly variable genes with minimum dispersion of 0.5 were selected. Trajectory and pseudotime analysis were performed using SCANPY as described above. Neighbors were computed on the first 30 principal components setting the number of neighbors to 30 and a Force-Atlas graph was computed. Cells were clustered using Leiden setting the resolution to 1.4. Diffusion pseudotime was calculated and cell-to-cell connectivities were computed using PAGA and projected on FA layout.

#### Supplementary Figure Legends

**Supplementary Figure 1. Immunophenotypic and morphological characteristics of *ex vivo* erythropoiesis.** (a) Flow cytometry time-course showing the expression profiles of two erythroid-specific cell surface markers, CD117 (C-Kit) and CD235a (Glycophorin A) over 12 days of culture. (b) Hematoxylin-Eosin staining of cytospin slides along the *ex vivo* erythropoiesis.

**Supplementary Figure 2. Biological replicates are highly concordant.** (a) Boxplots showing the correlation between donors and between days in DNase I-seq and RNA-seq experiments, respectively. (b) Scatterplots of log10 normalized Hotspot (DHS) counts from day 0 and $R^2$ between donors. Points are binned values and are colored by the number of points in each bin. (c) Scatterplots of log10 normalized gene counts (FPKM) from day 0 and $R^2$ between donors. Points are binned values and are colored by the number of points in each bin.

**Supplementary Figure 3. Differential enrichment for transcription factor binding motifs across DHS clusters.** Log$_2$-fold enrichment (x-axis) for binding motifs of known regulators of hematopoiesis across the DHS clusters (E10-E5, *y*-axis). Tripe asterisks indicates FDR < 0.05. N.S. indicates no significant enrichment.

**Supplementary Figure 4. Colocalization of developmentally co-regulated DHS and genes.** Clustering of TADs identified in CD34+ and day 11 erythroid Hi-C data, respectively, exhibiting enrichment for DHS or genes from specific clusters.

**Supplementary Figure 5. DHS clusters E2 and E3 display an extensive shared chromatin landscape with other myeloid cell types.** Percent of DHS in each cluster overlapping with DHS detected in $CD14^+$ monocytes and macrophages.

**Supplementary Figure 6. Erythroid specific DHSs are enriched for GWAS traits related to clinical erythroid phenotypes.** Frequency of GWAS SNPs associated with red blood cell (RBC) traits per Mb of all detected DHS (Hotspots FDR 5%), 11,805 developmentally regulated DHS and 5,816 late activated DHS from cluster E4 and E5.

**Supplementary Figure 7. Correlation based prediction of gene-enhancer connections identifies gene-proximal links.** Density histogram of genomic distances (Mb) of correlation-based predictions of enhancer-gene links (red) and chromatin loops predicted from day 11 *ex vivo* differentiated erythroid progenitor Hi-C data (blue).

**Supplementary Figure 8. Genetic knockouts confirm the predicted *cis*- regulators of *CDH1*.** (a) Position of TALE-FokI nuclease pairs (stylized as scissors) and primers (colored half-arrows) used against each of the upstream DHS and the promoter. (b) PCR-based validation of the genetic knockouts in each region. In HS2 (-12) an "in-out" PCR approach was followed. Wild type amplifies from outside forward primer (left yellow arrow) and inside reverse (green arrow). HS2 knock-out removes the binding site of the inside reverse primer and amplifies from the two outside primers, resulting in a larger fragment than wild-type.

**Supplementary Figure 9. Ranking TFs by average elastic-net regression coefficient per DHS highlights major *trans*- regulators during erythropoiesis.** Ranked TFs based on the average regression coefficient across each DHS cluster (E1-E5).

**Supplementary Figure 10. Assessing the capacity of elastic-net TF coefficients and raw TF motif instances per DHS to predict the DHS cluster.** Confusion matrices and overall error-rate of naïve Bayes classification of predicted DHS clusters using either discrete TF motif counts per DHS (top) or elastic net TF coefficients (bottom).

**Supplementary Figure 11. Enrichment of CTCF bound DHS in predicted loop anchors.** Density plot of DHS with > 1 CTCF motif in 5kb bins across a 300kb window around predicted loop anchors from *ex vivo* differentiated day 11 erythroid progenitor Hi-C data (black line) and permuted regions (grey line).

**Supplementary Figure 12. Assaying the clonogenic capacity during *ex vivo* erythroid differentiation.** (a) Schematic of the colony forming assay (MethoCult) during days 1-7 of *ex vivo* erythropoiesis along with representative microscope images of the detected types of colonies. (CFU-GEMM: Colony Forming Unit - Granulocyte/Erythroid/Macrophage/Megakaryocytic. CFU-GM: Colony Forming Unit - Granulocyte/Monocyte. BFU-E: Burst Forming Unit - Erythroid). (b) Changes in total clonogenic capacity during erythroid differentiation expressed as the number of colonies detected per 1,000 cells plated in methylcellulose assay. (c) Granulocytic/monocytic (CFU-GM) progenitor frequency per 1,000 cells plated during erythroid differentiation. (d) Cell expansion curve of the primary erythroid cultures from which cells were subjected to colony formation assays. The rate of growth is increasing monotonically over time.

**Supplementary Figure 13. A suspension-based assay to test for megakaryocytic potential during erythropoiesis.** (a) Schematic of megakaryocytic lineage potential assay during days 1-7 of *ex vivo* erythropoiesis where cells sampled daily were transferred to secondary suspension megakaryocytic media for 12 days. (b) Expression scatterplots of CD41a ($x$-axis) and CD42b ($y$-axis) of the cultures initiated from day 1 to day 7 of erythropoiesis. FACS profile of CD41 expression of a control megakaryocytic culture on day 12 is included as control.

**Supplementary Figure 14. *K*-means clustering of developmentally regulated genes during *ex vivo* megakaryocytic differentiation.** Dense gene expression time-course during *ex vivo* megakaryopoiesis and linear regression analysis identifies 5,487 significantly changing transcripts organized in 5 clusters (K1-K5) by *K*-means clustering.

**Supplementary Figure 15. Assessment of the erythroid potential along megakaryocytic differentiation in suspension assay.** Frequency of mature erythroid cells ($CD235a^+$) 12 days post transplantation into erythroid suspension culture of cells sampled from days 1-7 of primary megakaryocytic cultures.

**Supplementary Figure 16. Comparison of transcriptional dynamics along erythropoiesis and megakaryopoiesis between bulk and single-cell.** (a) Scatterplots of scRNA-seq gene expression (TPM) against total RNA-seq (FPKM) values and the respective $R^2$ values showing that data from the two experiments are highly concordant for erythrocytes (top) and megakaryocytes (bottom). Points are bins of individual genes where the color represents the number of points in a bin. (b) Representative examples of genes and their correlated expression between scRNA-seq and total RNA-seq experiments during erythropoiesis along with their Pearson correlation scores. (c) Representative examples of genes and their correlated expression between scRNA-seq and total RNA-seq experiments during megakaryopoiesis along with their Pearson correlation scores. (d) Hierarchical clustering of the samples using 10,000 highly variable, highly expressed genes displays data structure associated with sampling days and lineages.

**Supplementary Figure 17. Differential pseudotemporal dynamics between erythropoiesis and megakaryopoiesis.** Mean ± standard error of velocity pseudotime of single cells sampled along erythropoiesis and megakaryopoeisis.

**Supplementary Figure 18. Clusters of transcriptionally distinct cell states.** (a) Representation of Leiden clusters (0-17) and their PAGA connectivities (black lines) on Force-Atlas projection. Clusters are colored by average pseudotime. Cluster size (number of cells per cluster) is denoted as the size of circles. (b) Dendrogram of the Euclidean distances between Leiden clusters collapsed into biologically relevant populations (colored bars). (c) Topology of identified populations on the Force-Atlas projection. (d) The composition of populations as fraction of total cells sampled from each timepoint and lineage. (e) Heatmap of the expression (Normalized TPM) of representative erythroid, HSPC, megakaryocytic, and myeloid across populations. (f) Top significantly enriched gene sets found in the marker genes for each of the early populations (HSPC, MPP1, MPP2).

**Supplementary Figure 19. Identification of myeloid populations during *ex vivo* erythropoiesis and megakaryopoiesis.** Flow cytometry timecourse of the early myeloid marker CD33 (y-axis) during *ex vivo* erythroid (in red) and megakaryocytic (in blue) differentiation against CD117 (C-Kit) and CD41, respectively (x-axis).

**Supplementary Figure 20. Trajectory analysis and clustering of BM-derived hematopoietic progenitor populations.** (a) Diffusion pseudotime annotation of BM-derived hematopoietic populations on Force-Atlas projection. Solid arrows annotate the erythroid and myeloid trajectories. (b) Gene expression patterns of representative HSPC, Myeloid, Erythroid and Mk genes. (c) Leiden clusters (0-16) with PAGA connectivities (black lines) and average expression (log TPM) profiles per cluster for lineage marker gene sets. HSPC: *HOXA9, PROM1, THY1, FOS, CD34*. Myeloid: *SPI1, MPO, GATA2, CD33, CEBPA, CEBPB, IL3RA, CTSG*. Early Erythroid: *TFRC, KIT, CD36, CDH1, KLF1*. Late Erythroid: *HBB, HBA2, MXI1, ENG, ALAS2, GYPA*. Mk: *PLEK, ELF1, FLI1, TBP, ITGA2B, PPBP, PF4, GP9, MEIS1*.

**Supplementary Figure 21. Single-cell transcriptional dynamics of ex vivo erythropoiesis recapitulate erythroid population trajectories in the bone marrow.** Histogram of correlation (Spearman's $\rho$) of scRNA-seq expression profiles of the top 1,000 highly expressed genes between *ex vivo* erythroid differentiation and bone marrow erythroid trajectories (top left panel). Gene profiles for a selection of marker genes from *ex vivo* erythroid trajectories and bone marrow erythroid population trajectories. Trajectories are inferred from PAGA transitions between clusters or populations whereby cells within are ordered by pseudotime.

#### Supplementary Tables

**Supplementary Table 1.** A tab-separated table of the GRCh38 coordinates of all detected DHS (Hotspots FDR 5%) during *ex vivo* erythroid differentiation. mean is the average normalized DNase I density across 12 timepoints and 3 donors. l2fc is the log2 fold-change from the lowest daily density (3-donor average) to the highest observed. FDR is the false discovery rate of the likelihood ratio test between the explicit and the reduced linear regression model for DNase I density. Cluster is the *K*-means ($k$ = 5) cluster assignment. DHS with NA assigned cluster were not considered as developmentally changing DHS (see methods).

**Supplementary Table 2.** A table of all the developmentally regulated transcripts during *ex vivo* erythroid differentiation. mean_FPKM is the average normalized FPKM between day 0 and 12 and across 3 donors. l2fc is the log2 fold-change from the lowest daily FPKM (3-donor average) to the highest observed. FDR is the flase discovery rate of the likelihood ratio test between the explicit and the reduced linear regression model for gene expression. Cluster is the *K*-means ($k$ = 5) cluster assignment.

**Supplementary Table 3.** A table listing the links between DHS (GRCh38 coordinates) and genes, along with distance from TSS, Pearson correlation, correlation test *p*-value.

**Supplementary Table 4.** An Excel file including the TALEN pair name and their sequence as well as primers used to detect edits. Bases in lowercase represent the spacer region between the two half-TALENs.

**Supplementary Table 5.** A table of all the developmentally regulated transcripts during *ex vivo* megakaryopoiesis. mean_FPKM is the average normalized FPKM between day 0 and 12 and across 3 donors. l2fc is the $\log_2$ fold-change from the lowest daily FPKM (3-donor average) to the highest observed. FDR is the false discovery rate of the likelihood ratio test between the explicit and the reduced linear regression model for gene expression. Cluster is the *K*-means ($k$ = 5) cluster assignment.

**Supplementary Table 6.** Significant marker genes per single-cell population as identified by pairwise Wilcoxon-sum rank tests between populations. Test scores, p-values, adjusted p-values, log2 fold-change, and population contrasts are listed. Genes are filtered for adjusted $p$-value < $10^{-5}$ and absolute $\log_2$ fold-change > 1.

**Supplementary File 1.** A BED file with the TADs called at 10kb resolution from CD34 HSPC Hi-C data obtained from Misfud et al., 2015

**Supplementary File 2.** A BED file with the TADs called at 10kb resolution from day 11 *ex vivo* derived erythroid progenitors form Hi-C data obtained from Huang et al., 2017.

**Supplementary File 3.** A bedgraph file with Mustache loops calculated from TADs at 10kb resolution from Day 11 *ex vivo* derived erythroid progenitors.
