## Supplementary material for "Chromatin dynamics during hematopoiesis reveal discrete regulatory modules instructing differentiation": Supplemenal figures

### Supplementary Figure 1

**a**

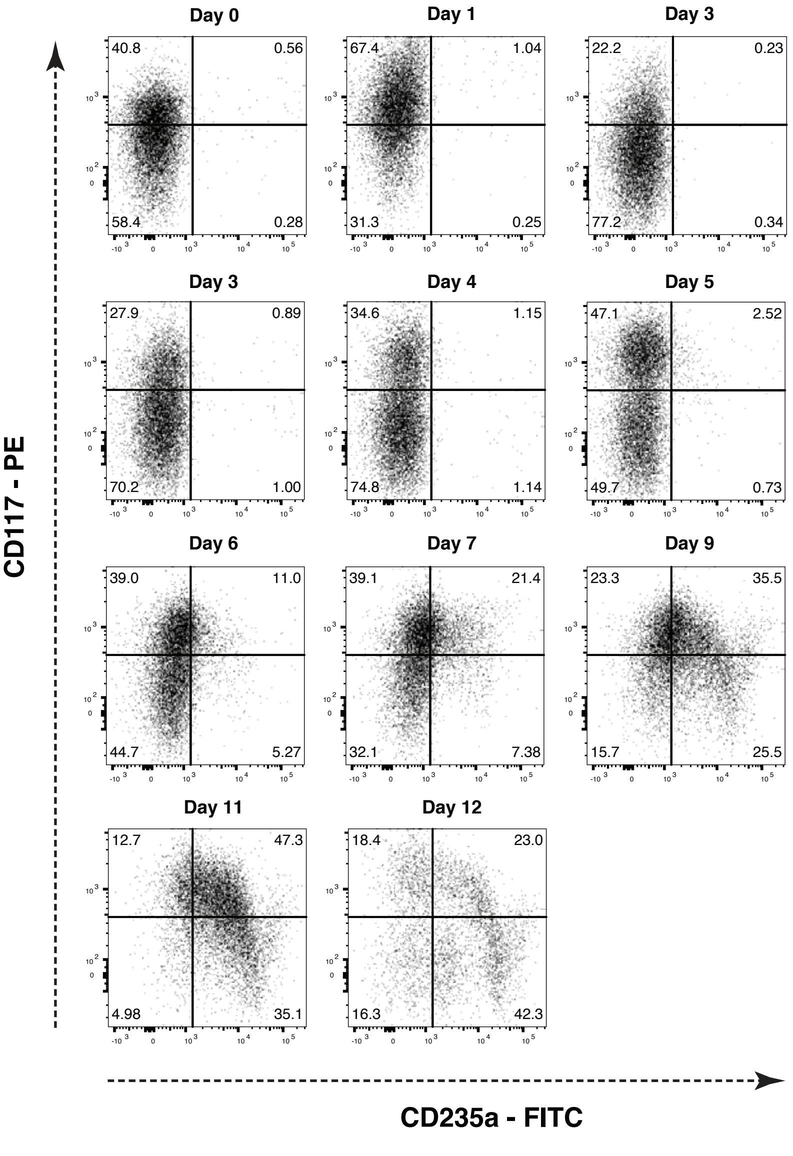


**b**

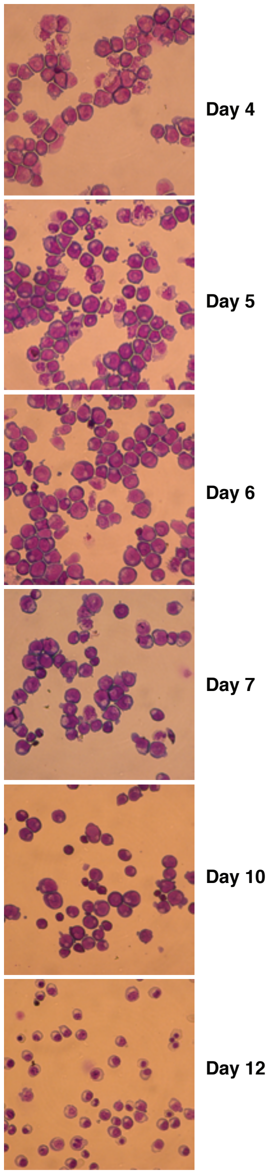

### Supplementary Figure 2

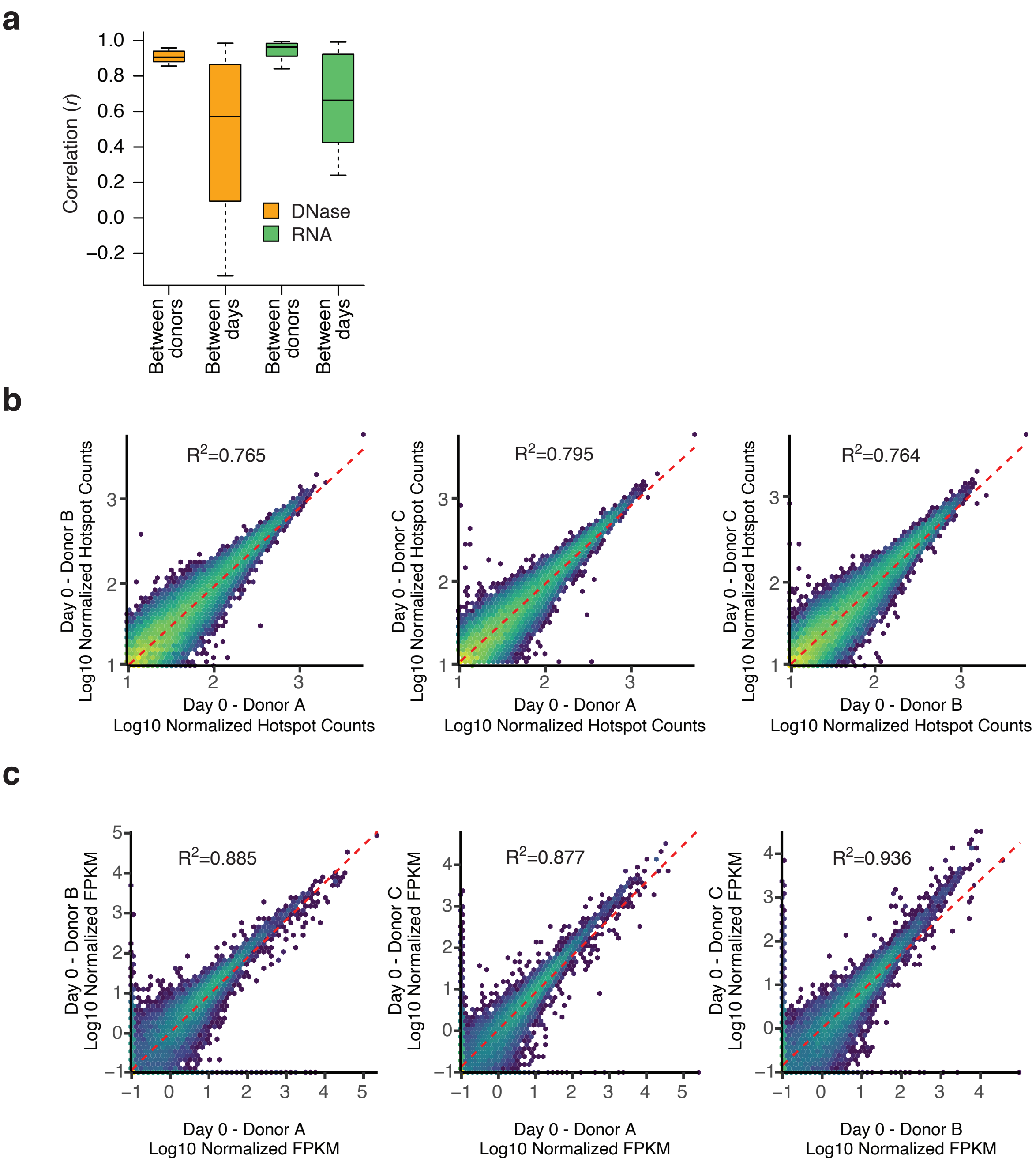

### Supplementary Figure 3

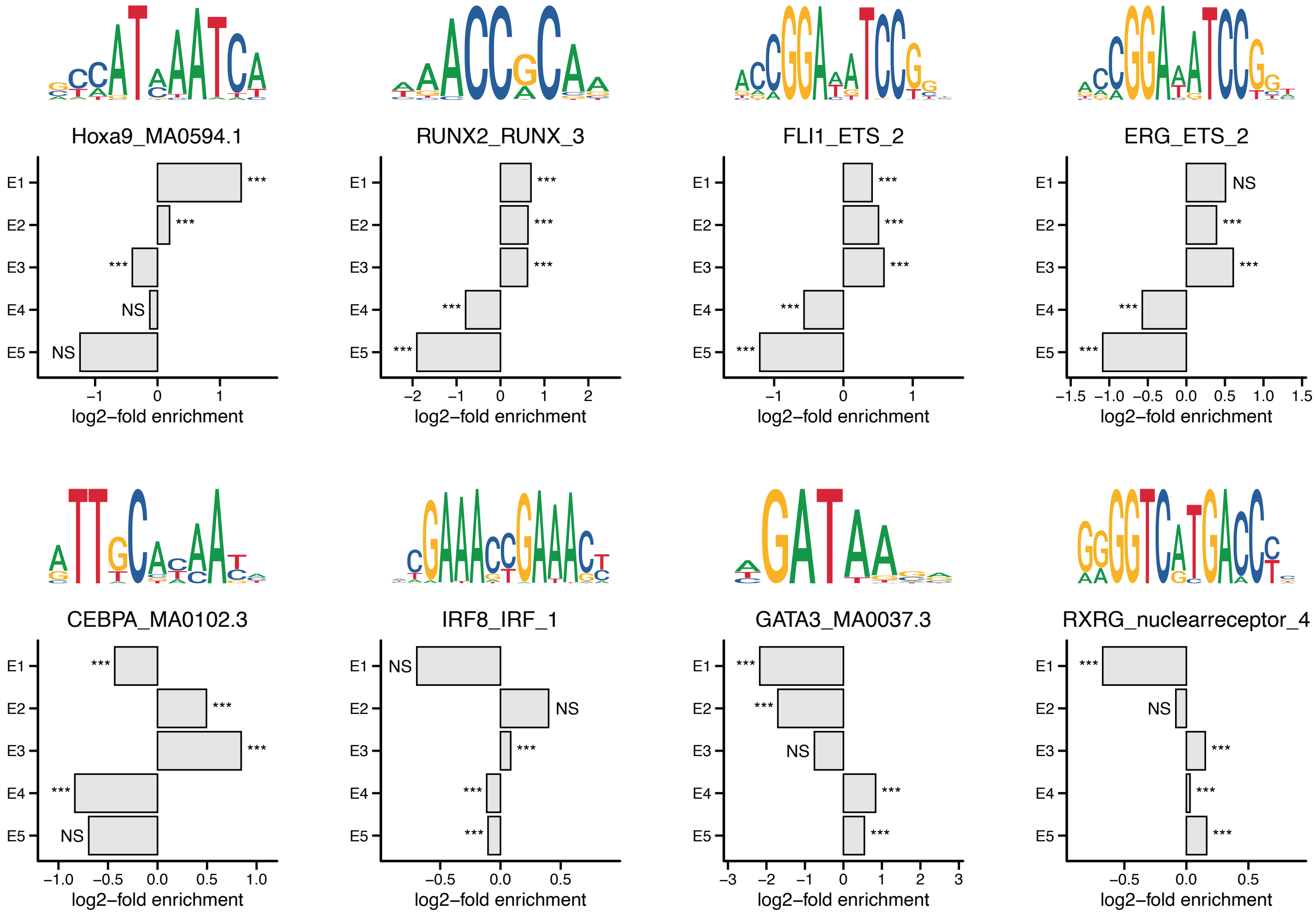

### Supplementary Figure 4

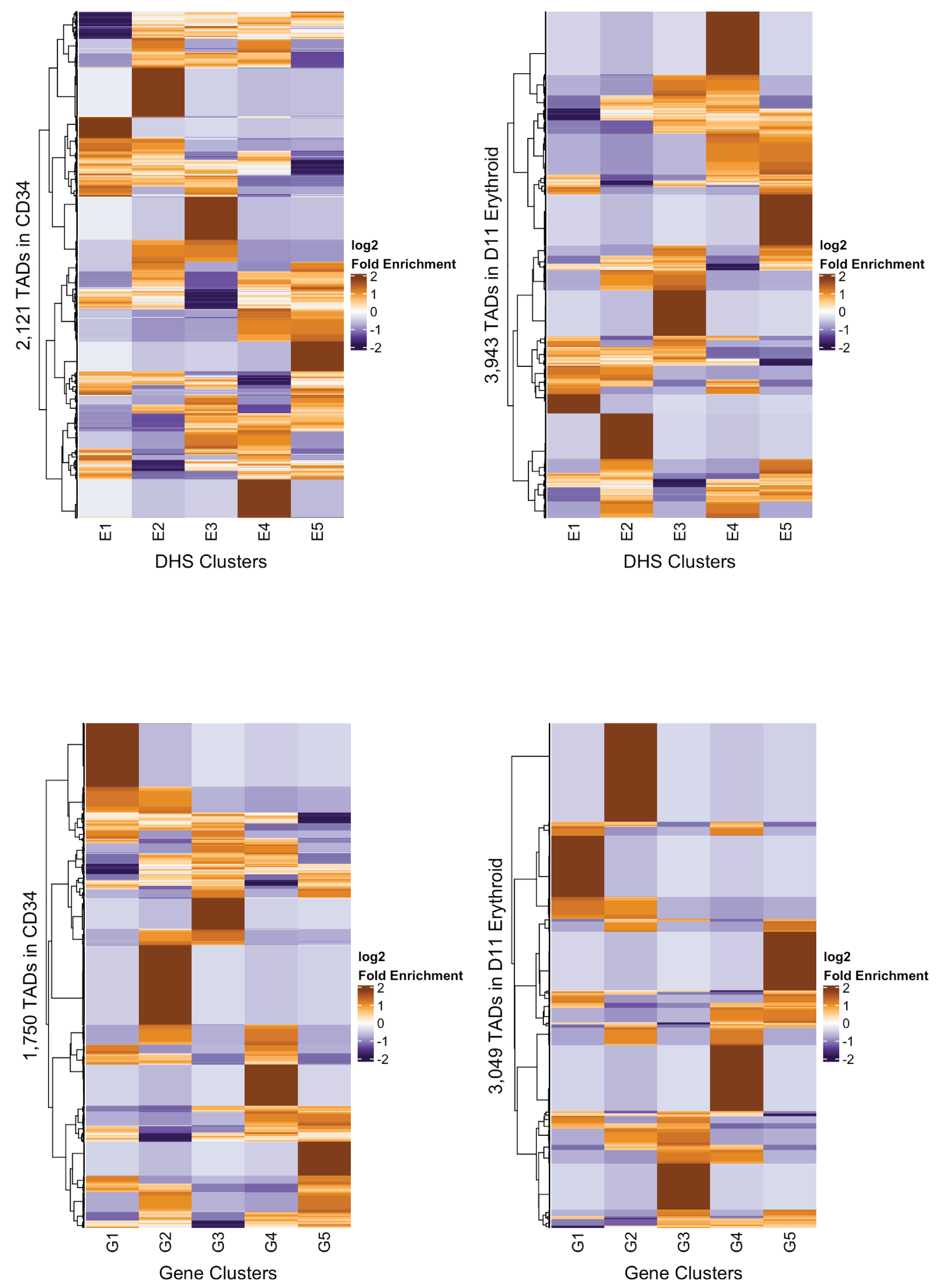

#### Supplementary Figure 5

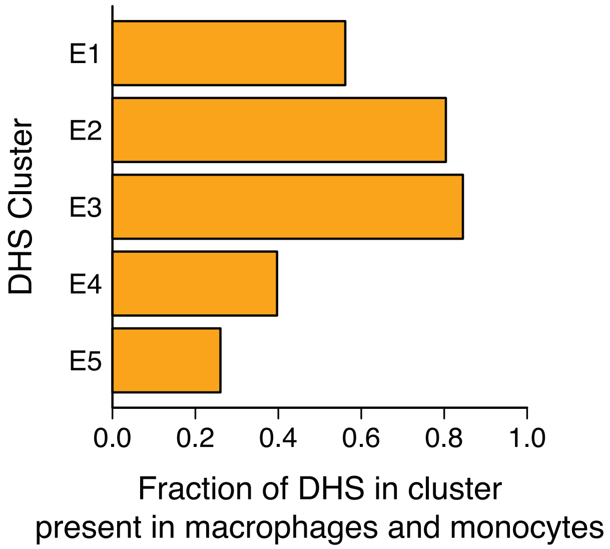

### Supplementary Figure 6

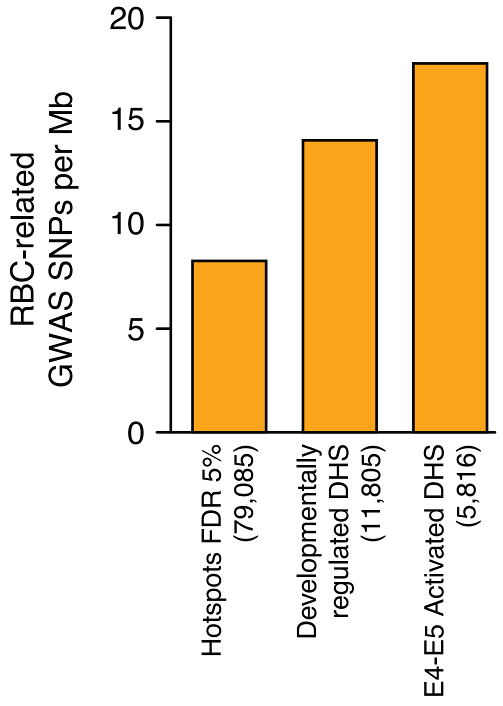

### Supplementary Figure 7

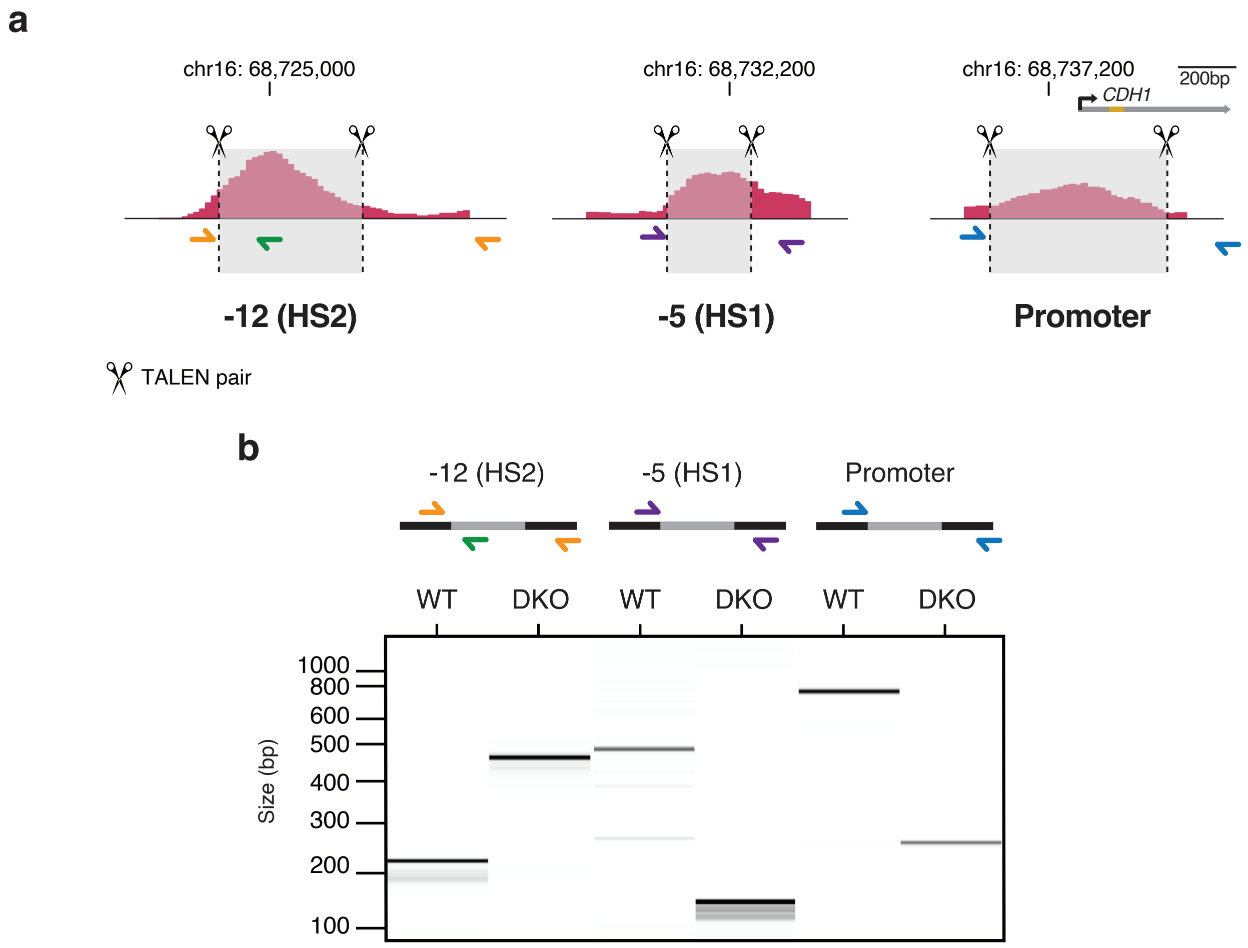

**Supplementary Figure 8**

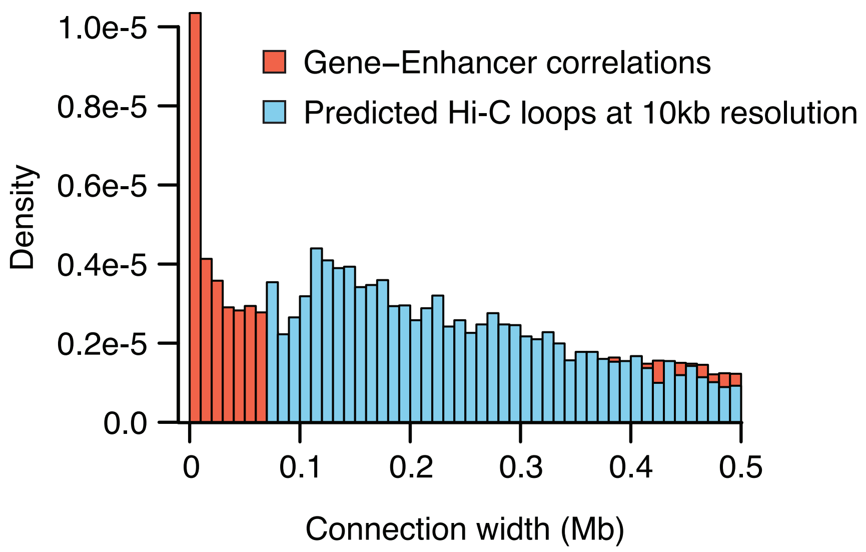

#### Supplementary Figure 9

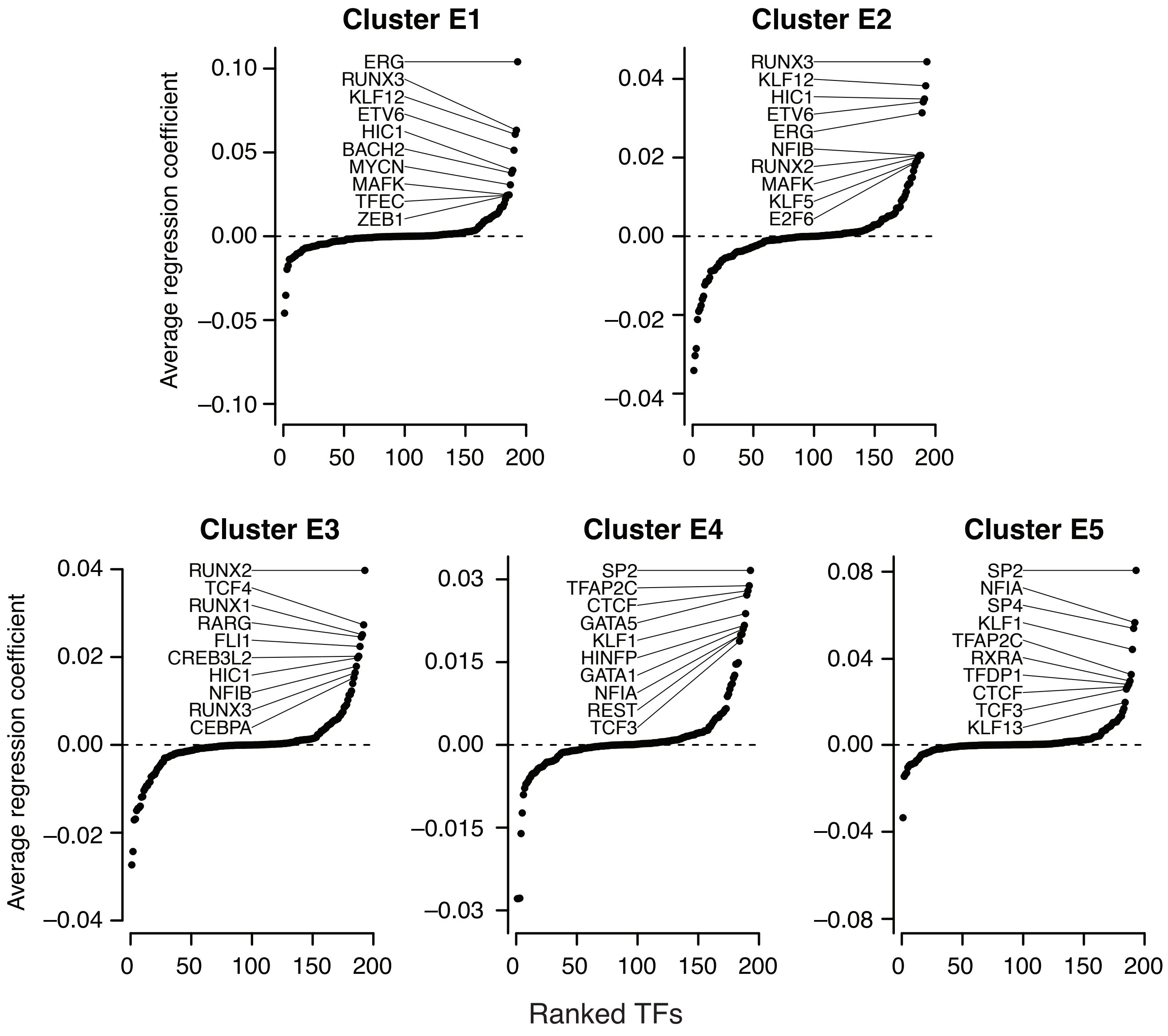

### Supplementary Figure 10

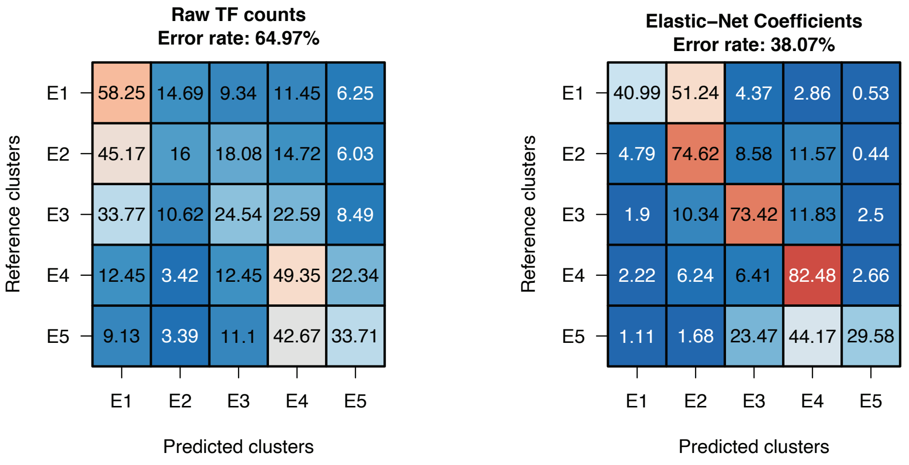

#### Supplementary Figure 11

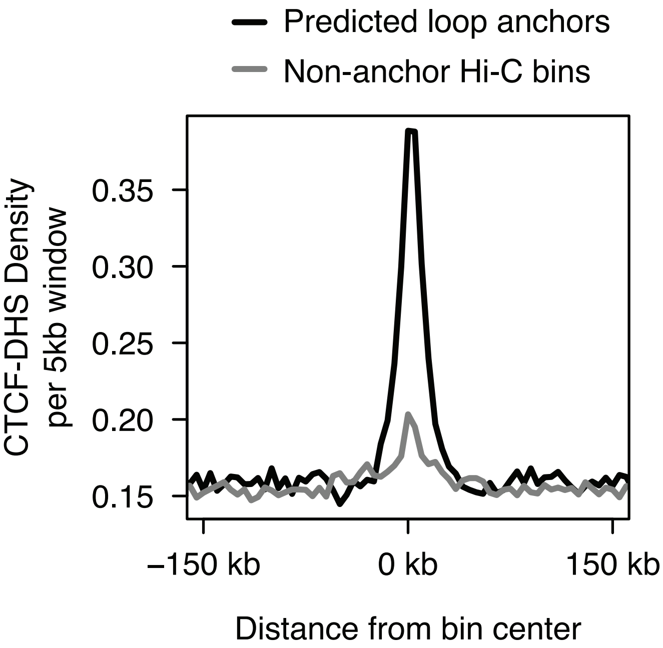

### Supplementary Figure 12

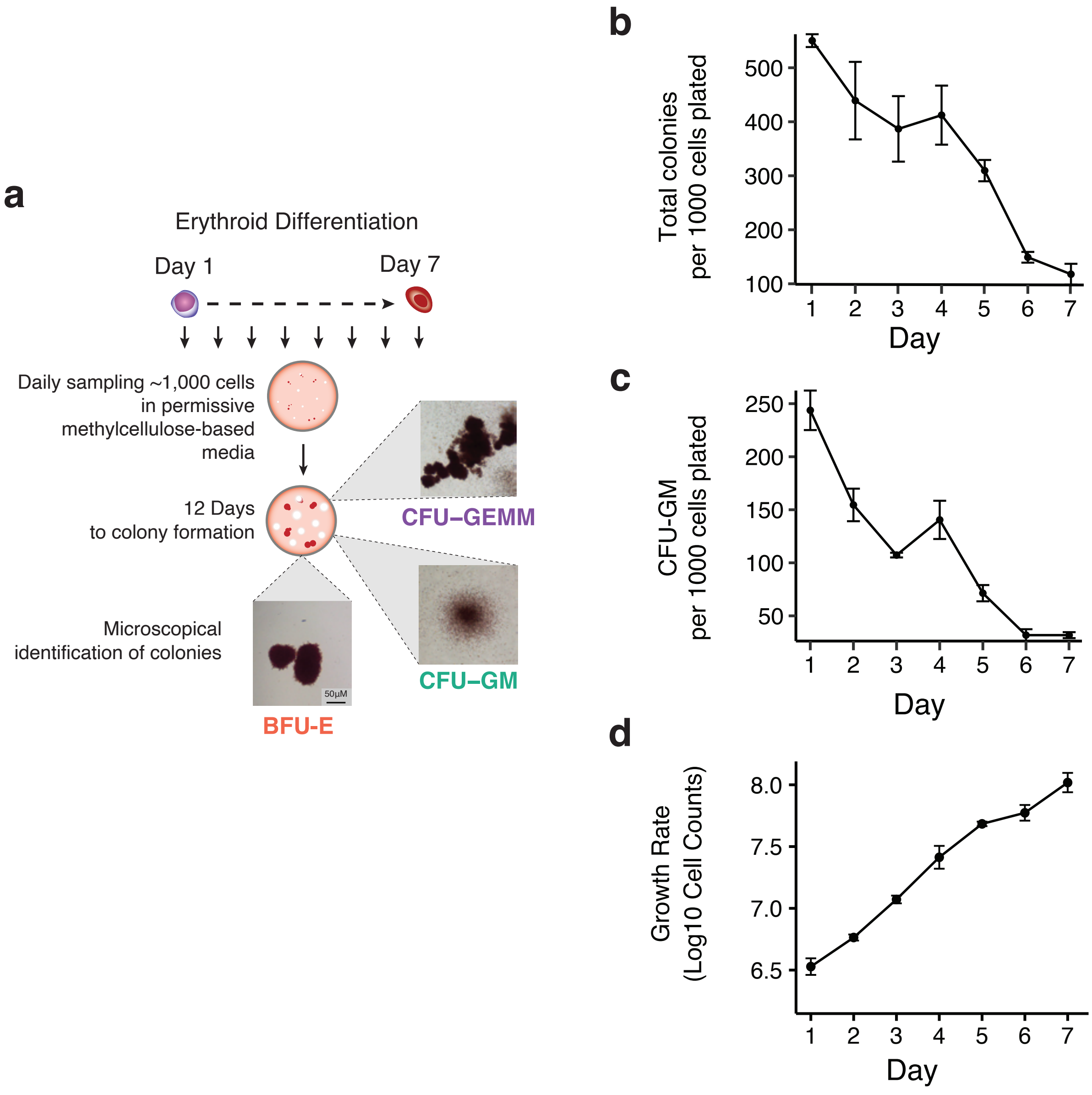

#### Supplementary Figure 13

**a**

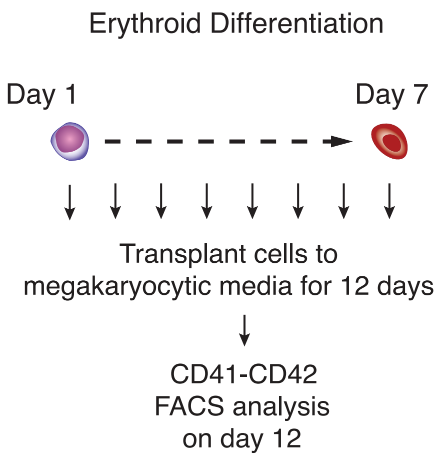


**b**

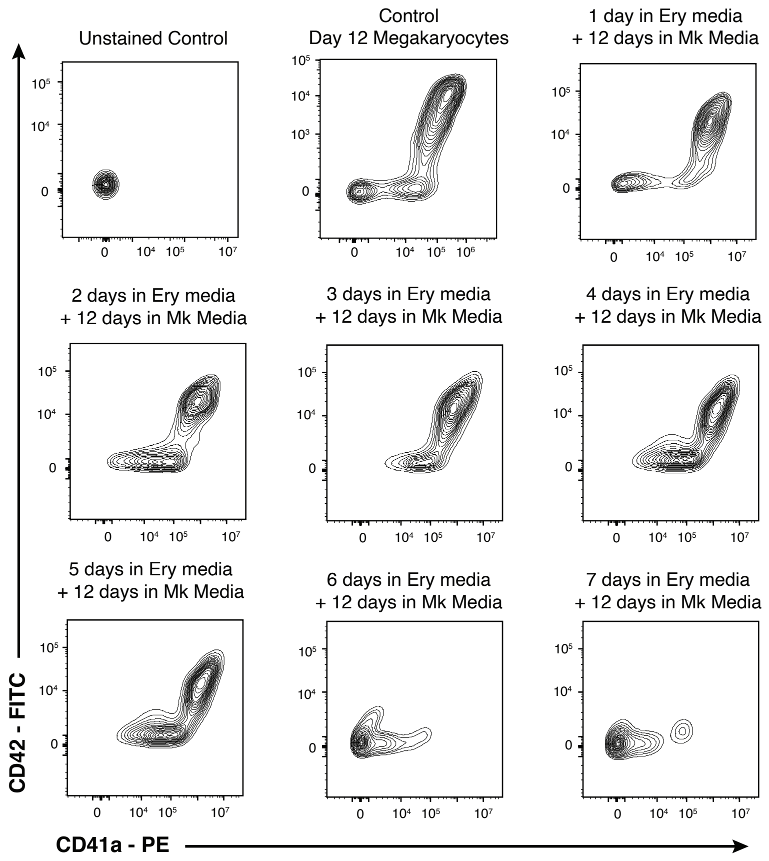

### Supplementary Figure 14

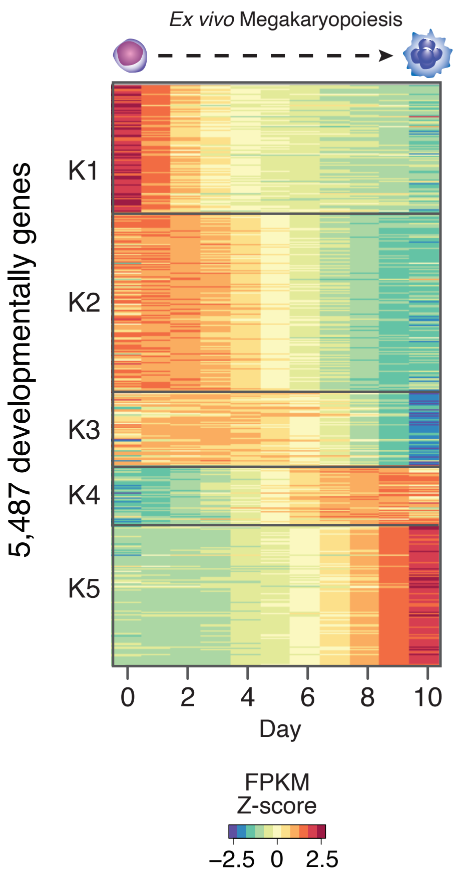

### Supplementary Figure 15

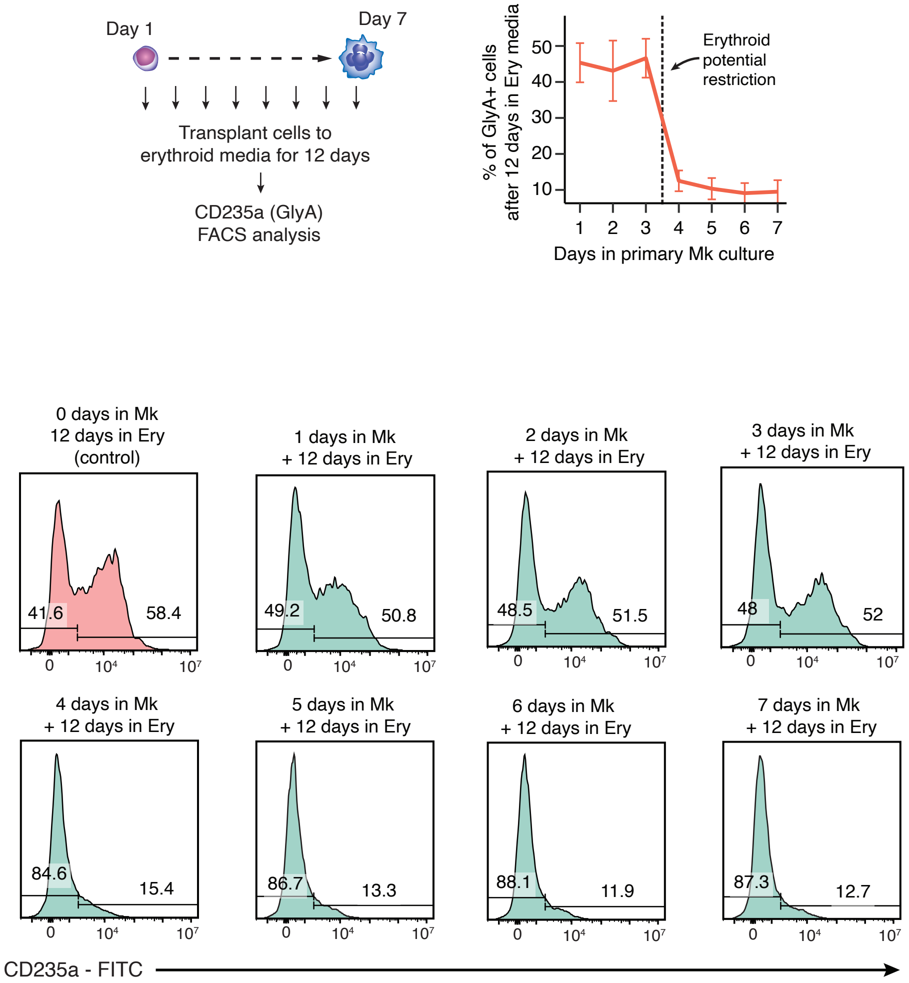

#### Supplementary Figure 16

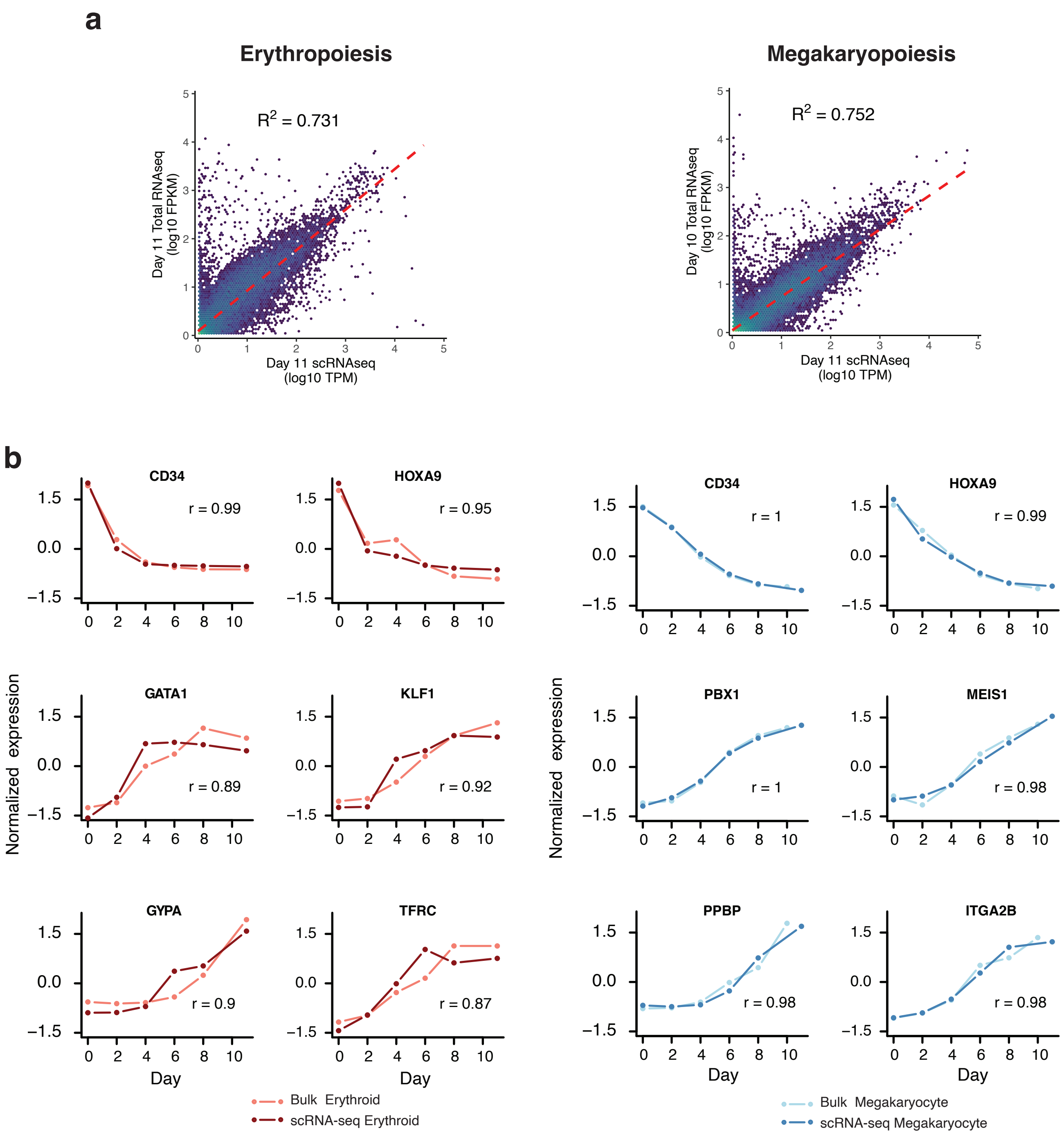

#### Supplementary Figure 17

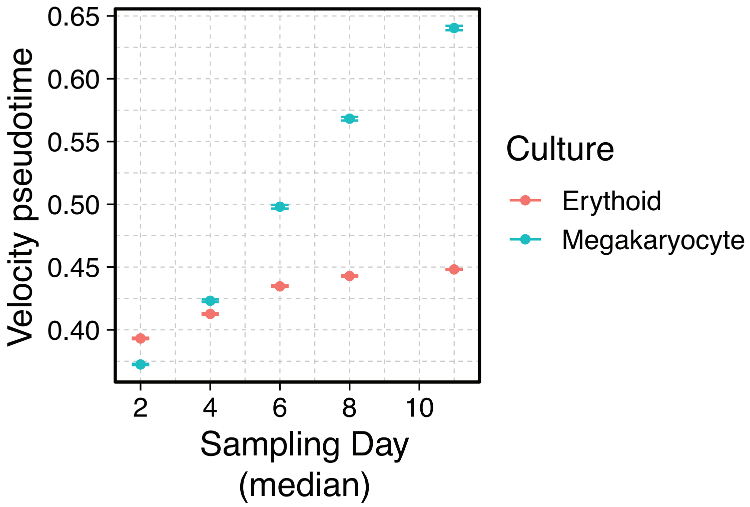

### Supplementary Figure 18

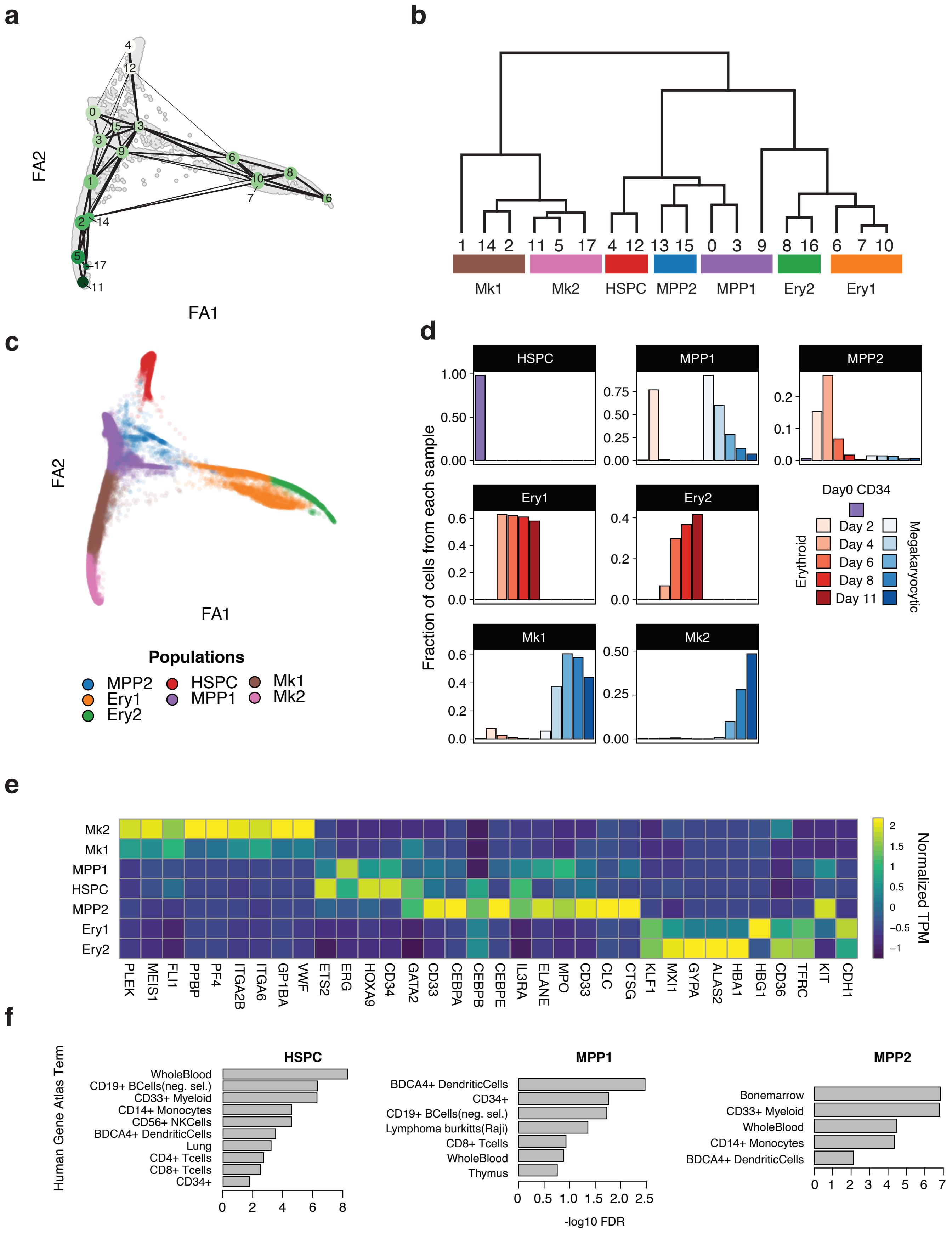

### Supplementary Figure 19

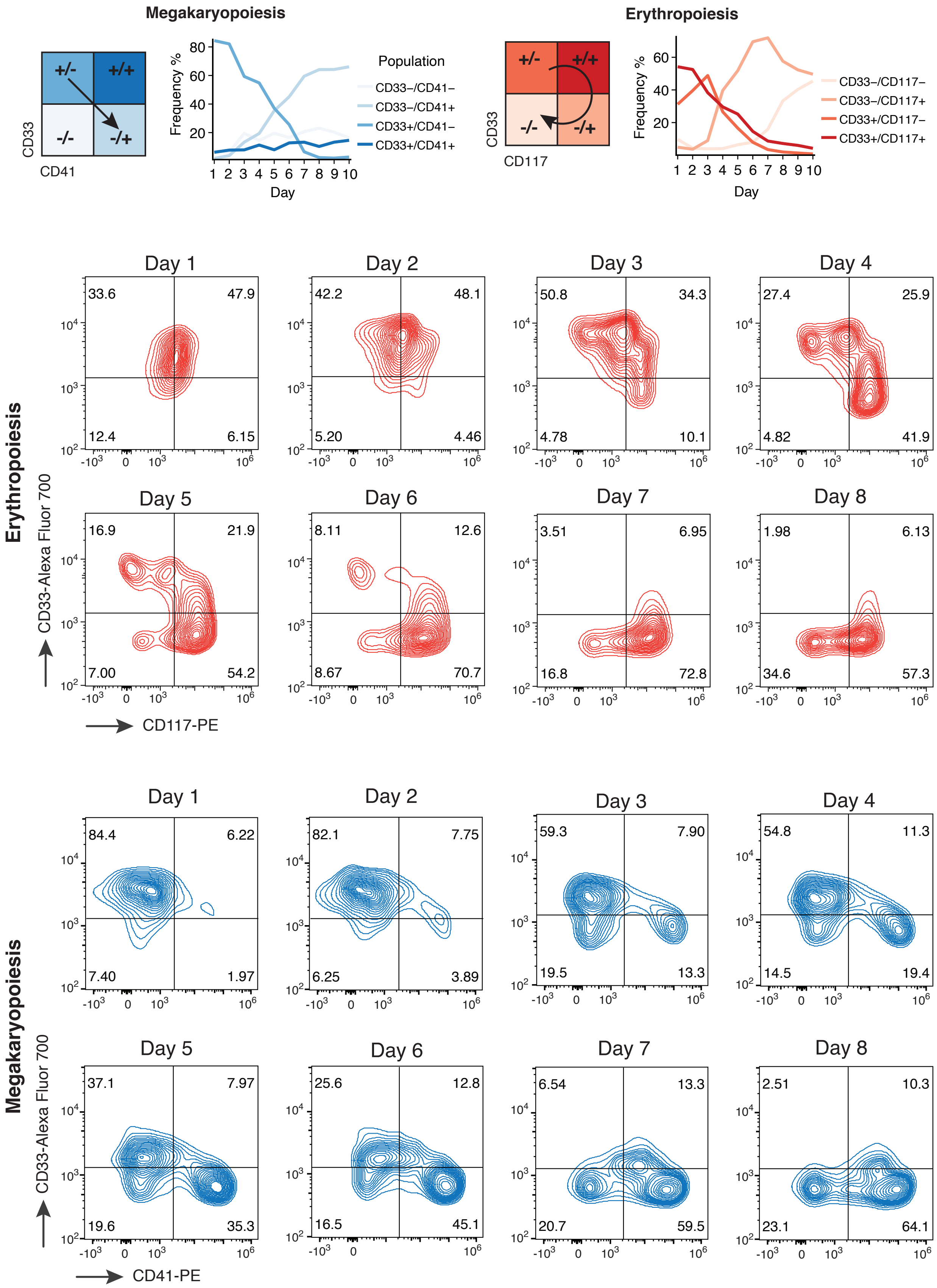

### Supplementary Figure 20

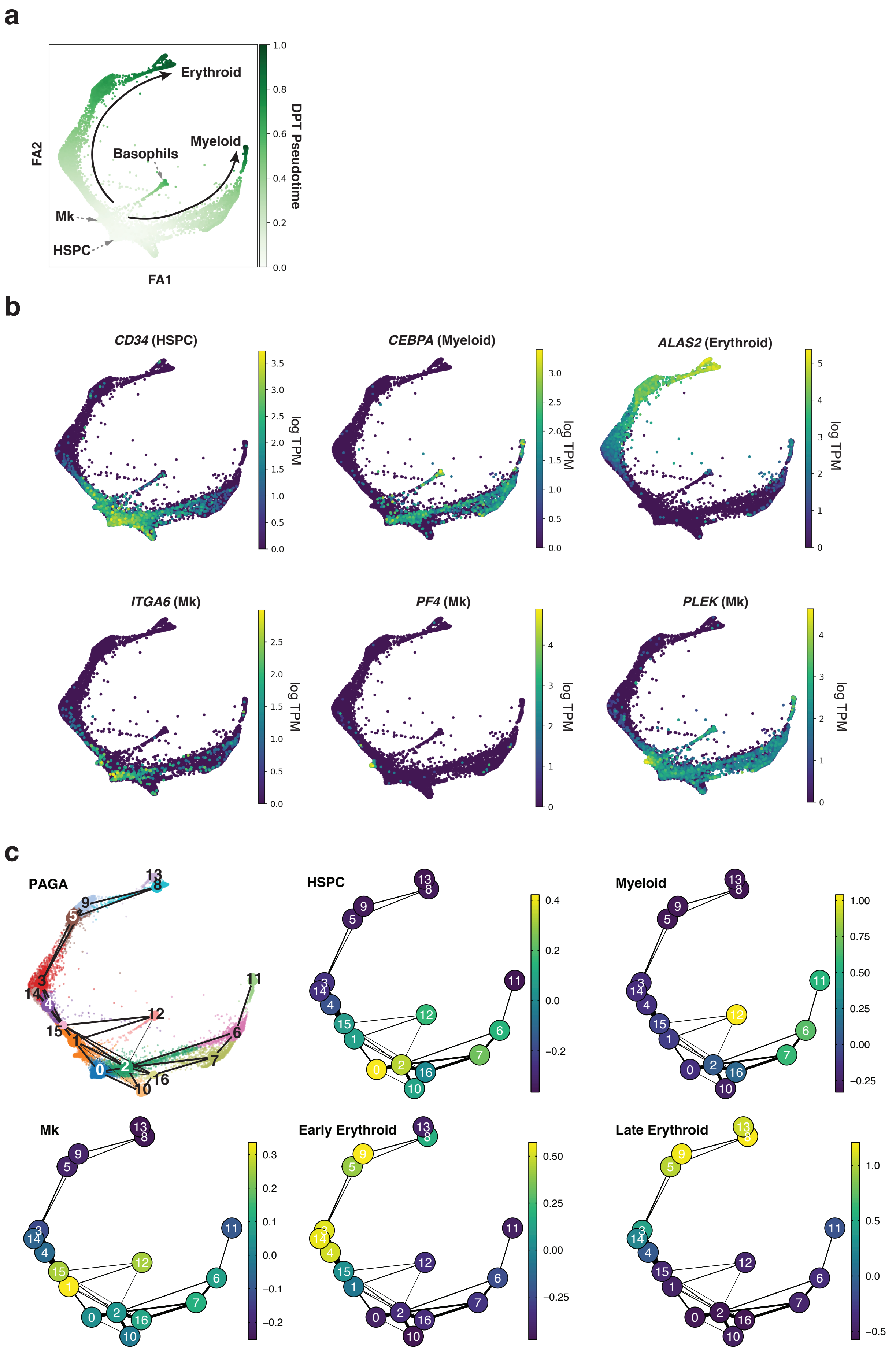

#### Supplementary Figure 21

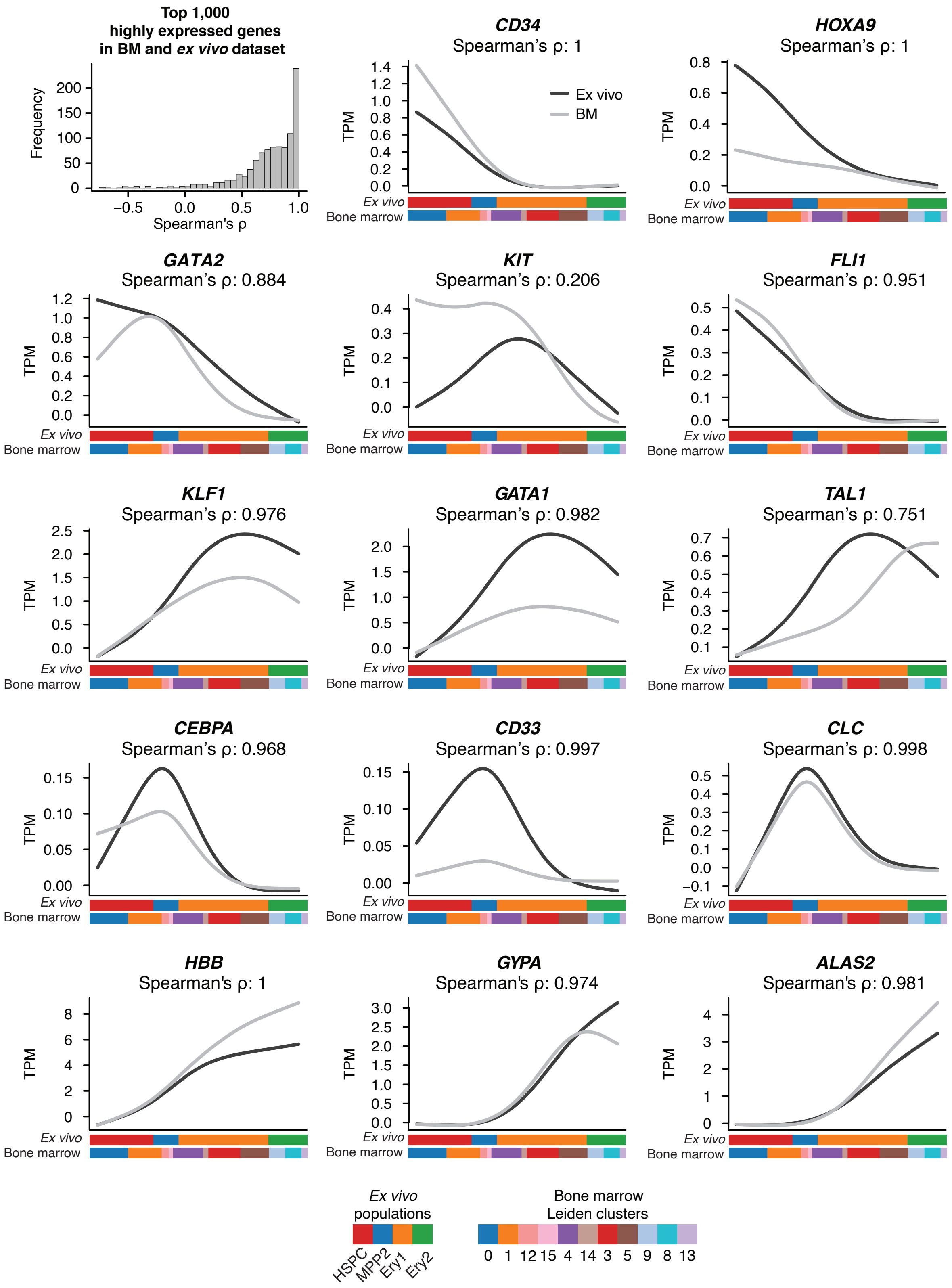
